# Integrative profiling of condensation-prone RNAs during early development

**DOI:** 10.1101/2024.10.14.618182

**Authors:** Tajda Klobučar, Jona Novljan, Ira A. Iosub, Boštjan Kokot, Iztok Urbančič, D. Marc Jones, Anob M. Chakrabarti, Nicholas M. Luscombe, Jernej Ule, Miha Modic

**Affiliations:** National Institute of Chemistry, Ljubljana, Slovenia; The Francis Crick Institute, London, UK; Dementia Research Institute at KCL, London, UK; Department of Basic and Clinical Neuroscience, Institute of Psychiatry, Psychology and Neuroscience, King’s College London, UK; PhD Program ’Biosciences’, Biotechnical Faculty, University of Ljubljana, Ljubljana, Slovenia; J. Stefan Institute, Ljubljana, Slovenia; University College London, UCL Respiratory, London, United Kingdom; Okinawa Institute of Science and Technology, Okinawa, Japan

**Author notes:** These authors contributed equally (Tajda Klobučar, Jona Novljan, Ira A. Iosub).

**Keywords:** Condensation-prone RNAs, semi-extractability, OOPS, RIC-seq, RNA-RNA interactions, deep learning, RNA-protein networks, phase separation, condensation

## Abstract

Complex RNA–protein networks play a pivotal role in the formation of many types of biomolecular condensates. How intrinsic RNA features contribute to condensate formation however remains unclear. Here, we integrate tailored transcriptomics assays to identify a distinct class of developmental condensation-prone RNAs termed ‘smOOPs’ (<u>s</u>e<u>m</u>i-extractable, <u>o</u>rthogonal <u>o</u>rganic <u>p</u>hase <u>s</u>eparation-enriched RNA<u>s</u>). These transcripts are localised to larger intracellular foci, form denser RNA-RNA interaction subnetworks than expected and are heavily bound by RNA binding proteins (RBPs). Using an explainable deep learning framework, we reveal that smOOPs harbor characteristic sequence composition with lower sequence complexity, increased intramolecular folding and specific RBP binding patterns. Intriguingly, these RNAs encode proteins bearing extensive intrinsically disordered regions and are markedly predicted to be involved in biomolecular condensates, indicating an interplay between RNA- and protein-based features in phase separation. This work advances our understanding of condensation-prone RNAs and provides a versatile resource to further investigate RNA-driven condensation principles.

## Introduction

Cells exhibit a wide range of RNA assemblies that physically partition them into subcellular membraneless compartments or biomolecular condensates, however the general molecular rules governing the RNA condensation and their local entrapment in ribonucleoproteins (RNPs) remain unclear^1,2^. RNA binding proteins (RBPs) and RNAs have both been implicated in condensate formation and disruptions in their phase separation have been linked to pathological conditions including impaired embryonic development, cancers, neurodegenerative diseases, and others^3–5^. Many proteins within ribonucleoprotein (RNP) condensates contain intrinsically disordered regions (IDRs) that are able to form weak multivalent interactions^6,7^) and simple changes to protein sequence or charge alone can drastically alter their condensation properties^8,9^. Conversely, RNA molecules can drive condensation themselves, either as a scaffold or through RNA-RNA interactions (RRIs)^10–15^. Recently, G3BP1 was shown to work as an “RNA condenser” that promotes intermolecular RRIs that stabilise stress granules^16,17^, while exceptionally long cytoplasmic mRNAs were shown to scaffold FXR1 protein into a network mediating signalling response^18^. Understanding which and how RNA features contribute to condensate formation and function, particularly through their interplay with RNA binding proteins (RBPs), remains a challenging question.

Several transcriptomic approaches opened the avenue for exploring condensation principles in an RNA-centric manner. Some of these methods are capturing RNAs based on their biochemical properties and their associations with RNA-binding proteins (RBPs), key for recruitment into and stabilisation within condensates^19^. Studies on semi-extractable RNAs^20,21^ identified a diverse array of RNA species associated with biomolecular condensates, particularly within nuclear bodies^21^. UV crosslinking-based methods recover RBP-bound RNAs^21,22,23^. In addition, RNA proximity-ligation approaches (reviewed in^24^ furthered our understanding of higher-order RNA structures, e.g. in stress granules^25^, as well as intermolecular RRIs, such as those between enhancer RNAs/mRNAs^26^, snoRNA/target RNAs^25^, and viral/host transcripts^27^.

To advance our current understanding of RNA-centric condensation features, we designated a novel class of transcripts as ‘smOOPs’ due to their <u>s</u>e<u>m</u>i-extractability^20^ and pronounced affinity for <u>o</u>rthogonal <u>o</u>rganic <u>p</u>hase <u>s</u>eparation (OOPS)^22^. Together, these two methods provide a comprehensive strategy to identify candidate RNAs that may be involved in condensation processes—that is, **condensation-prone RNAs**, which we define as highly interacting RNA molecules that are likely to enrich and concentrate in phase-separated compartments or other RNP assemblies. This group includes RNAs known to form or localise within condensates, alongside other RNAs sharing similar properties, which we hypothesise are prone to condensation. Through RNA *in situ* conformation sequencing (RIC-seq)^26^, we show that smOOPs engage in intermolecular interactions with one another, suggesting these RNAs are closely associated within cells. Using a combinatorial deep learning approach, we identified the core smOOPs features and revealed that these transcripts code for highly disordered proteins with elevated phase-separation propensity. Notably, this study also offers a comprehensive methodological framework for uncovering the principles of RNA assemblies and their possible role in coordinating post-transcriptional gene regulation, accelerating the extraction of condensation-relevant features across diverse datasets.

## Results

### Atlas of semi-extractable and OOPS-enriched RNAs across early development

To specifically enrich RNA molecules within RNP assemblies, we employed a combination of the semi-extractability assay^20^ and the OOPS assay^22^ (**Figure 1A**). We applied these methods to identify such RNAs during three distinct timepoints of early embryonic development: naïve pluripotent stem cells (nPSC) in 2iLif conditions^28^, primed epiblast stem cells (pPSC), in which lineage priming was prevented by Wnt inhibition^29,30^, and the earliest Wnt-differentiated primitive streak progenitors (dPSC)^31^. Standard TRIzol RNA extraction protocol was used as control. We generated total RNA-seq libraries that showed high correlations in gene-level counts within each developmental stage and assay type (**Figure S1A**). In a principal component analysis (PCA), the samples separated primarily by developmental stage (76% of the variance), with nPSCs being more distinct from pPSCs and dPSC. The assay type further contributed to the separation (14% of the variance), with the OOPS samples being the most distinct (**Figure S1B**).

**Figure 1:**
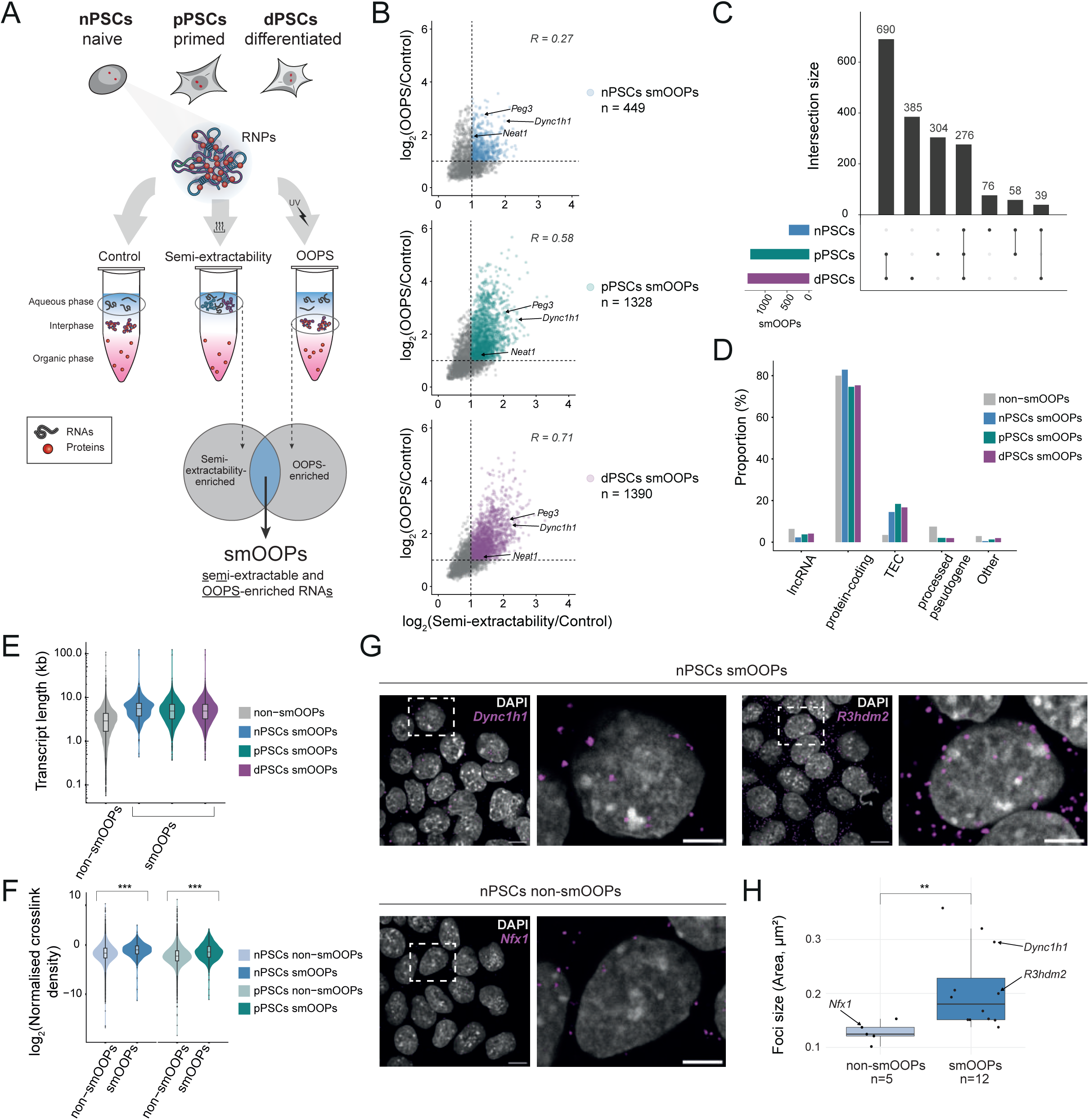
Atlas of semi-extractable and OOPS-enriched RNAs across early development. A) Experimental framework to identify RNAs that are both semi-extractable and highly RBP-bound RNAs (smOOPs) using three different TRIzol-based RNA extractions: aqueous phase of non-crosslinked sample as control, aqueous phase of heated and sheared TRIzol sample as semi-extractable RNAs and interphase of crosslinked sample to obtain OOPS-enriched RNAs. (nPSCs - naïve pluripotent embryonic stem cells (ESCs), pPSCs - primed pluripotent stem cells, dPSCs - 1day Wnt-differentiated pPSCs) B) Scatter plots showing the overlap between semi-extractability and OOPS-enriched genes (padj < 0.01) as well as the correlation between their fold changes compared to control at each developmental stage C) UpSet plot showing the intersections between smOOPs identified in nPSCs, pPSCs, and dPSCs. D) Percentage of gene biotypes for smOOPs and non-smOOPs in each cell-state. (TEC: To be Experimentally Confirmed). E) Comparison of transcript length distribution between smOOPs and non-smOOPs. F) Global iCLIP crosslinking signal normalised to expression and length and expression (CPM density/semi-extractability TPM) for smOOPs compared to non-smOOPs. G) Representative HCR-FISH photomicrographs (scale bar: 5 μm), with the right panel showing a magnified view of the region outlined by the dotted white box (scale bar: 10 μm). H) HCR-FISH quantifications. Boxplot showing the mean of foci size for each target transcript, calculated as the mean of all foci for each transcript. (Welch’s t-test, **p<0.01). n indicates the number of different mRNAs against which the HCR-FISH probes were designed. In total > 70 nuclei for a single transcript were imaged and > 890 foci counted for each transcript.

We next performed differential expression analysis to identify semi-extractable and OOPS-enriched genes compared to standard TRIzol RNA-seq controls at each developmental stage (**Figure S1C, Figure 1B, Data S1-S6**). By employing stringent effect-size and statistical cutoffs (at least two-fold enrichment and adjusted p-value < 0.01), we identified 449, 1328 and 1390 high-confidence genes at each developmental stage with distinctly increased semi-extractability and elevated RBP occupancy in OOPS samples, henceforth collectively referred to as smOOPs (<u>s</u>e<u>m</u>i-extractable and <u>OOPS</u>-enriched RNA<u>s</u>) (**Figure 1B,C, Figure S1C, Table S1**). Among smOOPs, we recovered RNAs known to form condensates, such as *Neat1* involved in paraspeckles^20,32^, *Dync1h1* known to form cytoplasmic foci of 3-7 copies at active translation sites in *Drosophila*^33^, and *Peg3,* for which the human homologue was found enriched in stress granules^34^. These examples underscore the inclusion of known condensate-forming RNAs within the smOOPs group (**Figure 1B**).

Of the total 1828 unique smOOPs, 276 were common to all stages, and most occurring at later stages. Notably, there were fewer unique smOOPs identified in nPSCs (76) compared to the other stages (pPSCs, 304; dPSCs, 385, **Figure 1C**). A positive correlation was observed between fold changes in the OOPS and semi-extractability assays, particularly upon the onset of cell fate commitment. However, the degree of enrichment (fold change) for a gene in one assay (OOPS or semi-extractability) did not always reflect the degree of enrichment in the other (**Figure 1B**). Some smOOPs highly enriched in OOPS were not similarly enriched in semi-extractability (**Figure S1D**), indicating that each assay preferentially captures distinct RNA characteristics, and that the smOOPs pool is rewired during developmental transitions (**Figure 1B,C, Figure S1D**). In terms of gene biotype, smOOPs predominantly consist of protein-coding genes (74.6%-82.2%), a small fraction of lncRNAs (2.2%-4.1%) and TEC genes (“To be Experimentally Confirmed”; 14.5%-16.7%) (**Figure 1D**). Notably, the TEC proportion was more than double that of the remaining genes not classified as smOOPs (non-smOOPs) (6.3%). The overrepresentation of TECs in the smOOPs group suggests that these understudied transcripts might play previously unrecognised roles in RNA-protein interactions or in the formation of RNP complexes. In conclusion, our approach highlights smOOPs as a group of transcripts both highly bound by RBPs and with unique extractability properties. This provides a developmental atlas of a class of RNAs that may have specialised roles in assembly of RNP complexes.

### smOOPs: condensation-prone RNAs form RNP granules

Our dual approach enabled us to identify smOOPs as candidate RNAs with potential for involvement in condensation processes. We observed that smOOPs are longer RNAs compared to non-smOOPs (**Figure 1E**), which may partially contribute to their distinct properties. To validate the tendency of smOOPs for RNP interactions, we performed global iCLIP - an orthogonal method for mapping the cumulative RBP occupancy across the transcriptome^35^ with nucleotide resolution. This not only confirmed the elevated RBP binding compared to non-smOOPs (normalised for expression and length; nPSCs p-value: 1.28×10^−1^^5^, pSCs p-value: 1.41×10^−4^^9^) (**Figure 1F**), but also provided precise positional information on RBP interactions (**Figure S1H**). We hypothesized that higher RBP occupancy in addition to the semi-extractability of smOOPs could indicate that they are part of RNP assemblies. To test this, we performed HCR-FISH^36^ using probes against the exons of 17 candidate protein-coding transcripts, including smOOPs and non-smOOPs (**Figure S1E**) with largely comparable expression levels (median TPM 21.9 for non-smOOPs transcripts and 32.5 for smOOPs; **Figure S1F**). Image analysis confirmed that smOOPs formed larger foci compared to non-smOOPs (**Figure 1G,H**) with higher overall intensity (**Figure S1G**), suggesting an enrichment of these RNAs in localised regions within the cell, potentially reflecting high local RNA concentrations. Notably, this pattern was also observed for smOOPs mRNAs that were already implicated in condensate formation*, Dync1h1* and *Peg3*^33,34^. Further analysis of RNA distribution found that only 36.9% of tested smOOPs foci were nuclear, compared to 46.5% for non-smOOPs transcripts (**Figure S1G**). Taken together, our findings suggest that smOOPs are a unique class of semi-extractable transcripts that are highly bound by RBPs and form larger foci within cells. These characteristics provide evidence of their condensation-prone nature.

### smOOPs establish RNA-RNA interaction subnetworks with enhanced connectivity

Given the well-established role of RBPs in regulating RNA assembly, we hypothesised that smOOPs participate in broader RNA networks, facilitated by RBP-mediated RNA-RNA contacts across the transcriptome. To globally map RBP-associated intra- and intermolecular RNA-RNA contact sites, we performed RNA *in situ* conformation sequencing (RIC-seq^26^) in nPSCs and pPSCs (**Figure 2A**). We detected 758,135 hybrid reads in nPSCs and 1,245,548 in pPSCs (excluding rRNA, tRNA, and mitochondrial reads), with 28% being intermolecular in nPSCs and 44% in pPSCs (**Figure 2B**). Gene-level intermolecular hybrid frequency was highly reproducible across samples (**Figure S2A**), with PCA showing developmental stage explained 88% of the variance (**Figure S2B**). The global distribution of contacts remained highly consistent across stages, with most involving intronic regions (∼66%, **Figure S2C**), similar to previous findings^26^.

**Figure 2:**
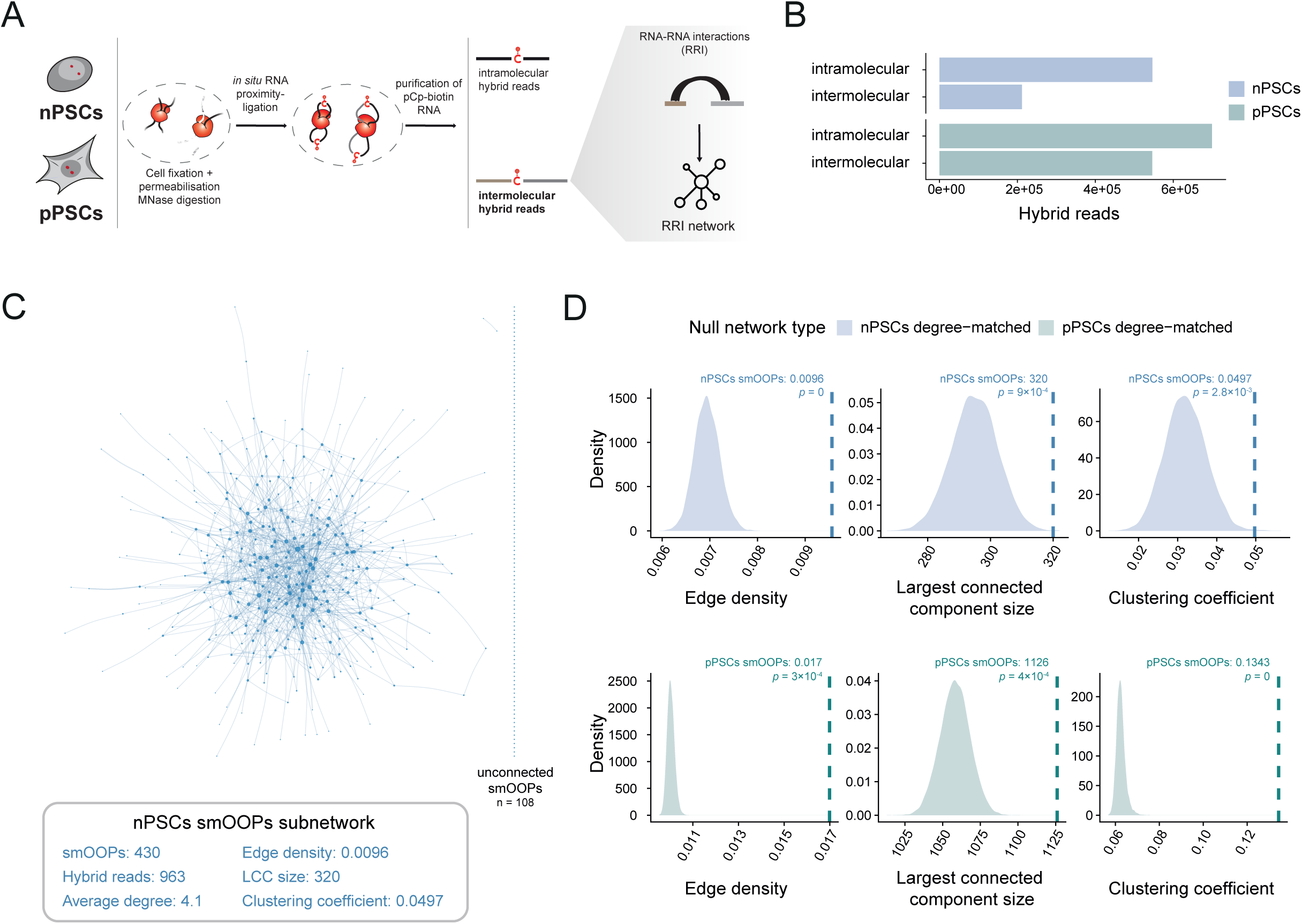
smOOPs’s connectivity within RRI networks. A) Schematic overview of the approach for inferring RNA-RNA interaction (RRI) networks in nPSCs and pPSCs using RIC-seq^26^, with key steps shown. B) RIC-seq hybrid read counts in nPSCs and pPSCs (pooled from three replicates each), categorised by interaction type, excluding hybrid reads containing rRNA, tRNA, and mitochondrial RNA. C) Visualization of the nPSCs smOOPs RRI subnetwork from RIC-seq data, where nodes represent genes and edges represent interactions based on hybrid reads. Node size corresponds to degree, and edge width represents the number of hybrid reads between nodes. Unconnected smOOPs are displayed on the right, and connectivity metrics below. D) smOOPs subnetwork connectivity comparison with degree-matched control subnetworks in nPSCs and pPSCs Density plots show the distribution of connectivity metrics from 10,000 degree-matched sub-sampled networks (null models). The dashed lines indicate observed values for the smOOPs RIC-seq subnetworks, with p-values from permutation tests comparing smOOPs subnetworks’ metrics to the metrics distributions for the degree-matched sub-sampled networks.

We generated developmental stage-specific RNA-RNA interaction (RRI) networks using intermolecular hybrid reads from RIC-seq, with genes as nodes and edges representing the frequency of interactions between them. The inferred networks showed typical characteristics of biological networks, such as protein-protein interaction (PPI) and RRI networks^37^. Specifically, the networks displayed scale-free-like behaviour, with most nodes having few connections, and a small number of highly connected nodes that dominate (**Figure S2D**). This suggests that a few genes play central roles in the network, while most genes have fewer connections, creating a heavy-tailed distribution of connectivity (**Figure S2D**). The RIC-seq networks also have small-world properties, with a higher global clustering coefficient than random networks of the same size, suggesting modular organisation (**Figure S2E**), and a relatively short average path length of 3.5, indicating that most RNAs are closely linked (**Figure S2F**).

Next, we explored the characteristics and connectivity of smOOPs in these RRI networks. In the nPSCs network, 430 of 449 smOOPs were present, and in pPSCs, 1255 out of 1328 smOOPs. In the RIC-seq networks, degree (number of distinct connections per gene) strongly correlates with expression for genes of similar length, regardless of whether they are classified as smOOPs (**Figure S2G**), likely due to the higher sensitivity of RIC-seq for detecting interactions in abundant RNAs. Although smOOPs appear to have high degree compared to all non-smOOPs, non-smOOPs with matched expression levels and lengths (**Figure S2H**) exhibit similarly high degree (**Figure S2I**). This suggests that the observed elevated degree of smOOPs is primarily driven by their expression and length, rather than their smOOPs status.

To explore interaction patterns among smOOPs, we examined the RIC-seq network focusing only on smOOPs-smOOPs interactions (i.e. smOOPs subnetworks) (**Figure 2C,D**). The nPSCs and pPSCs smOOPs subnetworks appeared highly connected, as indicated by key metrics: edge density (the proportion of possible edges present), largest connected component (the size of the largest connected subnetwork) and global clustering coefficient (the tendency of nodes to form tightly connected groups) (**Figure 2C,D**). To determine if the smOOPs subnetwork connectivity is greater than expected by chance, we compared it to null network models generated using degree-matched sub-sampling (see Methods), which accounts for biases linked to degree distribution (and accounting for expression level and length). Given the high degree of smOOPs (**Figure S2I**), this ensured that the observed difference in connectivity reflects underlying biological patterns, not merely the tendency of high-degree nodes to form more connections. Compared to the degree-matched sampled networks, smOOPs subnetworks exhibited significantly greater connectivity across all metrics in nPSCs and pPSCs (**Figure 2D**), indicating that their interconnectivity cannot be fully explained by their degree alone. Furthermore, despite the identity of smOOPs varying across development (**Figure 1C**), this characteristic is maintained in both nPSCs and pPSCs (**Figure 2D**). Together, these findings suggest that smOOPs are more interconnected among themselves, reflecting a specific network organisation that points to their proximity in cells.

### Deep learning accurately predicts smOOPs from intrinsic and regulatory RNA features

Given smOOPs are enriched in the semi-extractability and OOPS assays (**Figure 1**), and form denser RRI subnetworks than expected by chance (**Figure 2**), we pursued an in-depth investigation of the RNA features that define this distinct group of condensation-prone transcripts. Using intrinsic RNA features - such as sequence and structure - and transcriptome-wide data for trans-acting factors, we developed an explainable deep learning (DL) approach to distinguish smOOPs from a background RNA population that are neither semi-extractable nor OOPS-enriched. We focused on nPSCs due to the greater availability of public transcriptomic data compared to pPSCs, allowing us to utilise a more extensive set of features. Thus, for our binary classification, smOOPs from nPSCs with processed transcript length under 20 kb were used as the positive class (n = 447 out of 449), while genes without strong evidence of enrichment in either semi-extractable assay or OOPS vs control (padj > 0.01 and |LFC| < 1.4, see Methods) were defined as control (n = 1232) **(Table S1)**. The control genes have similar expression levels to smOOPs **(Figure S3A**), which strengthens our comparison by reducing potential confounding effects from expression differences. To identify the unique features of smOOPs, we trained DL classifiers with multiple feature sets: RNA nucleotide sequence, global RBP occupancy (global iCLIP, this study), m^6^A modification sites (miCLIP,^38^, transcriptome-wide base pairing intramolecular (PARIS-Intra) and intermolecular interactions (PARIS-Inter)^39^, *in silico* structure prediction with RNAfold^40^ and RNA-binding sites of 46 RBPs determined via CLIP from various mouse cell lines from the POSTAR3 database^41^. We encoded all features as positional information layers for each transcript, enabling efficient feature extraction (**Figure 3A**). To capture complex patterns and positional dependencies in the data, we implemented a DL architecture consisting of multiple convolutional neural network blocks (CNNs), a recurrent neural network (RNNs) and a multi-layer perceptron (MLP) (**Figure 3A**). The main goal was not merely to achieve accurate predictions, but also to extract and understand the classifying information contributed by each dataset, both individually and in combination. To accomplish this, we trained a separate model on every subset of the seven datasets, resulting in 127 unique models, each trained in eight replicates (**Figure 3A, Table S2,** see Methods).

**Figure 3:**
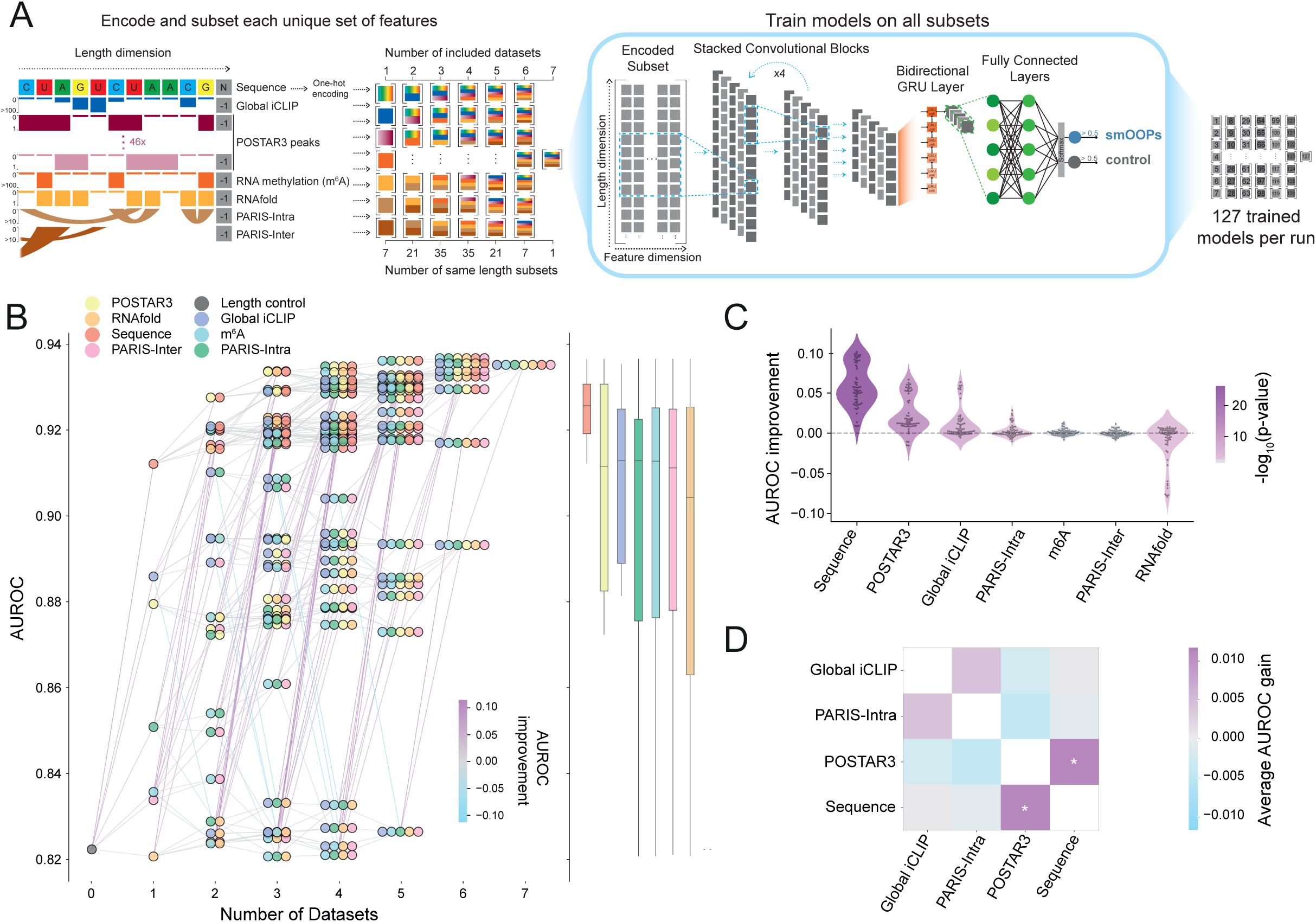
Deep learning-based classification of smOOPs and control transcripts. A) Schematic visualisation of the feature encoding and deep learning model training across 127 unique combinations of transcriptomic datasets. Each combination was used to train a model through convolutional and recurrent layers for nPSCs smOOPs and control transcript classification. B) Model performance across dataset combinations. (Left) AUROC of the best model trained on each unique dataset combination. Dots represent included datasets, with lines connecting models where one is a subset of the other, colour-coded by AUROC improvement. (Right) Boxplots showing the median performance and interquartile range as more datasets are included. C) AUROC improvement at the addition of a particular dataset to each combination of previously included datasets. Colour indicates the p-value from a two-sided Student’s t-test. D) Pairwise feature combination analysis quantifying the difference of maximum individual and combined contributions of features to model performance. An asterisk (*) marks statistically significant results (p-value < 0.05, two-sided Student’s t-test).

Given that smOOPs were longer than control transcripts (**Figure S3B**), and because all datasets implicitly encode transcript length due to padding, we additionally trained a baseline model using sequence length as the sole feature, also in 8 replicates **(Table S2)**. Transcript length alone achieves a good baseline performance (Area Under the Receiver Operating Characteristic Curve (AUROC) = 0.82, accuracy = 74%), which indicates that length provides meaningful information for smOOPs prediction (**Figure 3B**). The DL model trained only on sequence data greatly improved the prediction accuracy of the smOOPs not seen by the model (AUROC = 0.91, accuracy = 81%) (**Figure 3B, Figure S3C,D**), indicating that although length is important, sequence-specific features provide critical information for accurately distinguishing smOOPs. Notably, the sequence-based model outperformed models trained on any other individual dataset (**Figure 3B**, number of datasets = 1). The second highest predictive performance was achieved by models trained exclusively on global iCLIP data, with an AUROC of 0.89 and an accuracy of 79%. Since both iCLIP and OOPS rely on UV crosslinking to detect RBP binding, this high performance is likely due to an inherent overlap in methodological commonalities. Generally, as additional datasets were introduced, the predictive power gradually improved, ultimately reaching an AUROC of 0.94 and accuracy of 83% with all datasets included (**Figure 3B**, number of datasets = 7).

To highlight how each information layer contributed to the model’s predictive power beyond the others, we assessed the average AUROC improvement by comparing the model’s performance with and without each dataset. Our analysis revealed that the sequence, POSTAR3 peaks, global iCLIP, and PARIS-Intra layers significantly improved the performance of the models in which they were incorporated, giving us the confidence that these datasets contain important information about the features that distinguish smOOPs transcripts (**Figure 3C**). Surprisingly, RNAfold predictions reduced performance, presumably by introducing noise due to the limited ability of energy minimization algorithms to predict secondary structures for long sequences^42^ and the absence of context-specific information (e.g. cellular factors like RBPs or the influence of *in vivo* conditions that can affect folding). Most feature combinations showed minimal AUROC improvement, likely due to their multicollinearity (**Figure 3C**). To explore this overlap further, we analysed feature pairs from the top four most informative datasets (sequence, POSTAR3, global iCLIP, and PARIS-Intra) to assess their individual and combined contributions to AUROC. Excluding both features and adding one or both back revealed additive effects, particularly between RNA sequence and RBP occupancy. Notably, sequence data alone (which also reflects transcript length), encodes most information represented by the other datasets, except for individual RBP binding profiles, which likely refined the model by distinguishing sequences recognised by specific RBPs for better information extraction (**Figure 3D**). To validate the contribution of each feature across different transcript regions, we conducted masking experiments in which we omitted the 5’UTR, CDS, and 3’UTR signal (**Figure S3E**). We found that the sequence data contributed most to the predictive power in the CDS, while the global iCLIP and POSTAR3 were most informative in the 3’UTR. In contrast, the performance of the PARIS-Intra based models decreased when either the CDS or the 3’UTR was masked (**Figure S3E**).

Together, our analyses demonstrate that using the selected datasets within our DL framework enabled accurate smOOPs prediction, and that training models on all feature combinations revealed nuanced dataset interactions that might otherwise go unnoticed.

### Deconvolving the features of smOOPs

To identify the unique features distinguishing smOOPs, we used integrated gradients to compute nucleotide-resolution importance scores for the models trained on each of the top four datasets that improved the average model performance (i.e. sequence, POSTAR3, global iCLIP, and PARIS-Intra) . Overlaying the importance scores over each transcript uncovered distinct patterns of feature importance for each dataset (**Figure S3F**). We analysed global feature patterns by dividing smOOPs transcripts into 100 bins and averaging feature importance scores within each bin, capturing patterns across datasets. We then clustered smOOPs based on the importance scores across these feature dimensions, revealing two clusters (cluster 1, n = 328; cluster 2, n = 119) (**Figure 4A, Table S1**).

**Figure 4:**
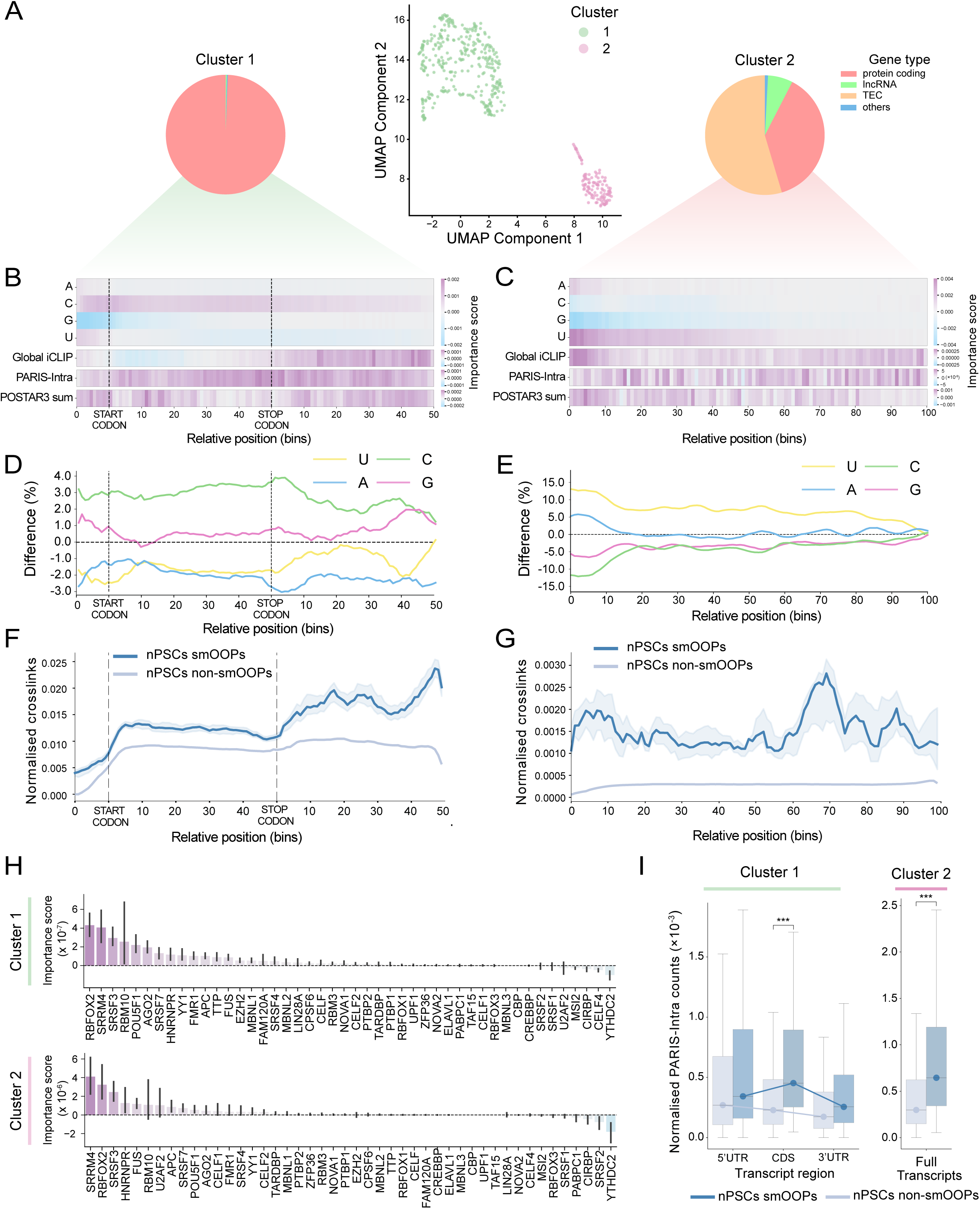
Analysis of smOOPs predictive features. A) UMAP projection of binned importance scores per feature for nPSC smOOPs. Each dot represents a transcript, colour-coded by cluster. The accompanying pie charts show the distribution of gene types within each cluster. (TEC: To be Experimentally Confirmed). B) Heatmap showing the average nucleotide- and dataset-specific feature importance scores for all transcripts in Cluster 1, divided into 10 bins for the 5’UTR and 50 bins each for the CDS and 3’UTR.. C) Heatmap showing the average nucleotide- and dataset-specific feature importance scores for all transcripts in Cluster 2, binned into 100 intervals along the transcript length.. D) Per-bin difference in average nucleotide content between cluster 1 smOOPs and control transcripts, with nucleotide content divided into 10 bins for the 5’UTR and 50 bins each for the CDS and 3’UTR. E) Per-bin difference in average nucleotide content between cluster 2 smOOPs and control transcripts, with nucleotide content divided into 100 bins across the transcripts. F) Median global iCLIP signal, normalised for expression and binned (10 bins for the 5’UTR and 50 bins each for the CDS and 3’UTR) for cluster 1 and control transcripts. Shaded areas represent 95% confidence intervals per bin, estimated via bootstrapping. G) Median global iCLIP signal, normalised for expression and binned (100 bins across the transcripts) for cluster 2 smOOPs and control transcripts. Shaded areas represent 95% confidence intervals per bin, estimated via bootstrapping. H) Average importance scores for CLIP datasets from POSTAR3 across all transcripts in clusters 1 and 2, shown for each RNA-binding protein with 95% confidence intervals.. I) Bar charts showing PARIS intramolecular hybrid counts across individual transcript regions for cluster 1 and smOOPs compared to all non-smoOPsl transcripts, adjusted for region length and expression (p-value < 0.001, two-sided Welch’s t-test).

Cluster 1 primarily contained mRNAs (98.2%), while cluster 2 was a mix of TECs (56.4%), mRNAs (36.4%) and lncRNAs (7.3%) (**Figure 4A**). We observed distinct sequence compositions in each cluster: cluster 1 exhibited a pronounced cytosine (C) enrichment across the transcript, particularly within the 5’UTR and CDS, which overlapped with high PARIS-Intra importance (**Figure 4B**). In contrast, cluster 2 displayed increased overall importance of uridine (U), with adenine (A) being slightly more important at the 5’ ends of the transcripts (**Figure 4C**). Although global RBP occupancy (global iCLIP) is a key feature across all smOOPs, our analysis highlighted positional specificity of RBP binding in each cluster: the importance of RBP occupancy was predominantly concentrated in the 3’UTR for cluster 1, while for cluster 2, it was distributed more uniformly across the entire transcript (**Figure 4B,C**). Furthermore, the regions of POSTAR3 importance, summing individual CLIP binding profiles (POSTAR3 sum), portrayed more specific regions of importance, emphasising the precise nature of RBP interactions at these sites (**Figure S4A,B**). Cumulatively, our DL approach pinpoints the key biological information for RNA condensation propensity, with sequence, intramolecular structure and RBP binding information being most predictive.

Comparing nPSCs smOOPs with all non-smOOPs transcripts, we sought to validate and deepen the insights gained from our models by directly investigating features highlighted as important. Building on model predictions, a lower sequence complexity compared to controls was confirmed for both clusters: the mRNAs in cluster 1 portrayed a C-rich CDS (**Figure 4D**), with the ‘CCC’ triplet being the most enriched, followed by ‘GCC’, ‘CGC’, ‘CCG’, and ‘CCA’ (**Figure S4C,E**). The most frequent triplets in the U-rich cluster 2 (**Figure 4E**) included ‘UUU’, followed by ‘UUA’, ‘UCU’, ‘UGU’, and ‘UAU’ (**Figure S4D,F**). Global iCLIP data showed slightly enhanced global RBP binding in the 3’UTR region of smOOPs in cluster 1 (**Figure 4F**), however there was a 10-fold greater median normalised RBP occupancy for transcripts in cluster 2 (**Figure 4G**). Since UV crosslinking preferentially targets uridines, we cannot determine the exact extent to which this bias influences the observed enrichment in RBP occupancy. Importance scores for POSTAR3 data revealed that in both clusters, RBFOX2, SRRM4 and SRSF3 carried key predictive information (**Figure 4H**). Since SRRM4 is not expressed at sufficient levels in our cell line, the model likely determined its importance based on its binding motifs. The frequency of intramolecular interactions determined by PARIS was also significantly increased in the CDS of smOOPs from cluster 1 and overall higher in smOOPs from cluster 2 (**Figure 4I**). Despite these specific features of smOOPs, we did not observe any major differences in translation efficiency^43^ or in mRNA stability^44^ compared to non-smOOPs (**Figure S4G-I**).

These findings highlight that smOOPs are more strongly bound by RBPs, generally more structured than non-smOOPs, and can be divided into two clusters with distinct sequence composition in nPSCs: one comprising C-rich mRNAs and the other predominantly A/U-rich transcripts. In both clusters RBP binding plays a crucial role, emphasising their role in shaping the behaviour of these unique transcripts.

### smOOPs mRNAs encode proteins rich in intrinsically disordered regions

Since smOOPs in nPSCs were clearly distinguished from control transcripts based on their sequence features, we investigated whether these features persisted throughout development. To address this, we trained an additional deep learning model using only the sequence of smOOPs transcripts identified at all three developmental stages - nPSCs, pPSCs, and dPSCs - and compared them to the same set of control transcripts **(Table S1)**. This newly trained model achieved performance comparable to that of the nPSCs-only model (AUROC = 0.90, accuracy = 82%), indicating that sequence features uniquely define smOOPs throughout the embryonic development, not only in nPSCs. Furthermore, when examining the model’s learned features, we again identified two smOOPs clusters—C-rich and A/U-rich—which showed 96% consistency with those from the nPSCs-only model **(Figure S5A,B, Table S1)**. This confirms that this separation is a general feature of smOOPs that persists throughout development .

Interestingly, the ratio of C-rich to A/U-rich smOOPs shifts during development. In nPSCs, C-rich smOOPs were predominant, comprising approximately 75% of nPSCs-specific smOOPs. In dPSCs, their proportion declined to 56%, reflecting an increased representation of A/U-rich smOOPs at later developmental stages (**Figure 5A, Figure S5C**). This observation prompted further investigation into how smOOPs behavior changes across semi-extractability and OOPS assays for the two clusters. While C-rich smOOPs maintained consistent enrichment across both assays, A/U-rich smOOPs became increasingly semi-extractable during development while showing reduced OOPS enrichment (**Figure 5B**). This coincided with a higher proportion of mRNAs in the A/U-rich cluster and a gradual decline of TECs (**Figure S5D**).

**Figure 5:**
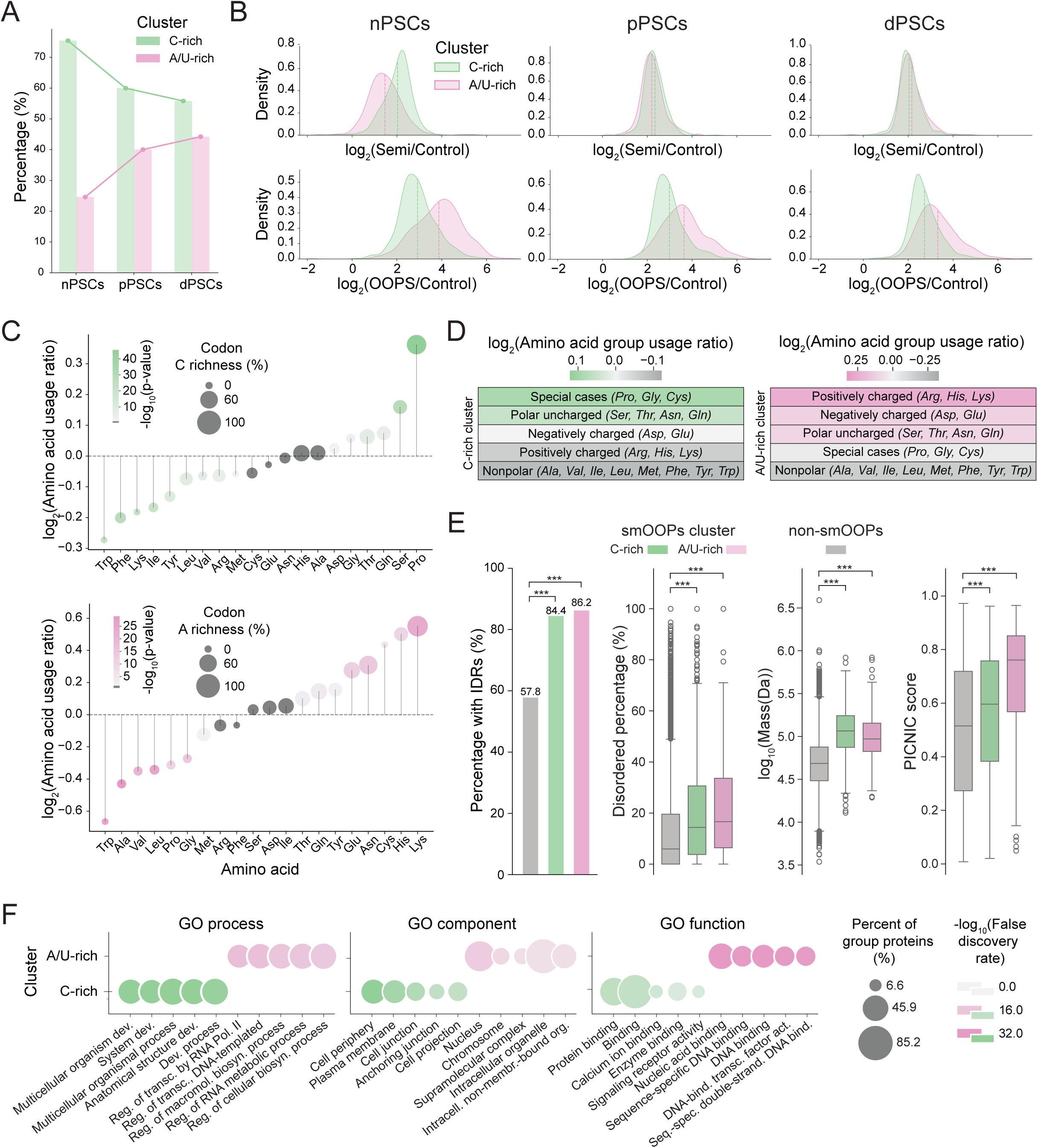
IDR content, amino acid composition, and functional characteristics of smOOPs-encoded proteins. A) Percentage of smOOPs transcripts in the C-rich and A/U-rich clusters across developmental stages (nPSCs, pPSCs, and dPSCs). B) Density distributions of semi-extractability and OOPS enrichment for smOOPs transcripts in the C-rich and A/U-rich clusters across nPSCs, pPSCs, and dPSCs. C) Amino acid enrichment in C-rich and A/U-rich smOOPs clusters compared to non-smOOPs. Dot size indicates the percentage of C/A nucleotides in codons encoding each amino acid, with gradient strength showing statistical significance. D) Enrichment of amino acid groups in C-rich and A/U-rich smOOPs clusters compared to non-smOOPs. E) Percentage of proteins with IDRs and measures of disorder for C-rich and A/U-rich smOOPs clusters compared to non-smOOPs. Bar chart shows the percentage of proteins with IDRs and the violin plots show the percentage of disorder in proteins, their mass and PICNIC score (Proteins Involved in CoNdensates In Cells)^46^. F) GO term enrichment analysis for C-rich and A/U-rich smOOPs clusters. The dot size represents the percentage of proteins enriched for each term, and the color intensity reflects statistical significance (FDR).

To elucidate the basis of the nucleotide composition differences between the smOOPs clusters, we examined the contribution of codon bias or amino acid composition. In line with the A/U-rich smOOPs nucleotide bias, the amino acid composition of this group was heavily enriched in charged and polar residues (**Figure 5C,D**). In contrast, C-rich smOOPs encoded proteins enriched in proline and polar uncharged residues, particularly serine (**Figure 5C,D**). Since many enriched amino acids (proline and serine in C-rich cluster, glutamine, glutamate and lysine in A/U-rich cluster) are all highly abundant in intrinsically disordered protein regions (IDRs)^45^, we investigated whether smOOPs encode proteins with an inherent propensity for condensation.

Structural analysis confirmed this hypothesis, revealing that both C-rich and A/U-rich clusters encode more proteins with IDRs and a higher percentage of disordered amino acids (**Figure 5E**). Notably, as many as 84.4% of C-rich and 86.2% of A/U-rich smOOPs-encoded proteins contain IDRs, while only 57.8% non-smOOPs-encoded proteins (**Figure 5E**). In addition, smOOPs-encoded proteins are, on average, more than twice the size of the non-smOOPs group, resulting in significantly longer IDRs. Both the number of IDR-containing proteins and and the proportion of disorder within these proteins (**Figure 5E**) suggested smOOPs-encoded proteins might be involved in condensate formation. We tested this using the PICNIC (Proteins Involved in CoNdensates In Cells) prediction model^46^, providing further evidence that smOOPs-encoded proteins are more likely to be involved in condensates (**Figure 5E, Figure S5E**). Interestingly, despite their shared structural properties and condensation potential, the two clusters encode proteins with distinct cellular functions. Gene ontology analysis revealed that C-rich cluster proteins are localised to the cell membrane and periphery, are mainly involved in developmental processes and play roles in protein binding. In contrast, A/U-rich cluster proteins are mostly nuclear, involved in gene regulation and contain nucleic acid-binding domains, especially zinc-finger motifs (**Figure 5F, Figure S5F, Table S3**).

These findings suggest that smOOPs encode two classes of proteins with high condensation propensity but distinct cellular roles. Differences at both the RNA and protein levels underscore the sequence-driven and developmentally regulated nature of smOOPs, highlighting their involvement in condensate formation throughout development.

## Discussion

Understanding the principles governing RNA assemblies in their native cellular context is crucial to uncovering how these structures shape cellular biology. In this study we use a dual methodology — semi-extractability assays and OOPS — to define smOOPs based on their shared biochemical properties during early murine embryonic development. This combination provides a comprehensive view of RNAs potentially involved in RNP assemblies and condensation. smOOPs are a distinct class of long transcripts that consists of predominantly protein-coding RNAs exhibiting subcellular localisation in larger foci. RIC-seq analysis furthermore reveals that smOOPs form more densely connected RRI networks than expected, altogether suggesting their potential spatial organisation and cellular compartmentalisation.

We systematically investigate their characteristics using an integrative DL approach that identifies two clusters based on sequence composition, RBP binding patterns and structuredness: C-rich mRNAs with structured regions and extensive 3’UTR RBP binding, and A/U-rich transcripts with high overall RBP occupancy. This heterogeneity suggests that smOOPs may contribute to condensation through diverse mechanisms or in different cellular contexts. Across the three developmental timepoints, transcripts in the C-rich cluster show stable enrichment in both methods, while those in the A/U-rich cluster become increasingly semi-extractable and less OOPS-enriched. This suggests that A/U-rich smOOPs undergo greater developmental changes in RNP assembly, and may play distinct roles in condensate formation at later developmental stages.

The most striking finding is the link between smOOPs’ RNA sequence features and the presence of intrinsically disordered regions (IDRs) in their encoded proteins. Disordered protein regions tend to be encoded by repetitive nucleotides or sequence motifs, such that the sequence repetitiveness is reinforced by codon biases^47^. Here we show that both smOOPs clusters encode a significantly higher proportion of proteins with IDRs compared to non-smOOPs. However, the two clusters are distinguished by differences in nucleotide and amino acid composition, which likely contribute to the distinct functionalities of their encoded proteins. This finding hints at a possible coordination between RNA identity and the phase separation potential of the proteins they encode, a concept that requires further systematic investigation.

While earlier studies have examined the roles of specific RNAs or RBPs in condensation, this study takes a broader approach by identifying and characterizing an entirely new class of RNAs with potential implications for phase separation. This resource and our findings provide a foundation for future research aimed at confirming the involvement of smOOPs in condensation, unraveling the functional relevance of the two identified clusters, and elucidating the mechanisms by which RNA features may coordinate with protein disorder and phase separation potential.

In terms of methodology, our study demonstrates the power of explainable machine learning to reveal complex patterns across diverse datasets, enabling unbiased classification and characterisation of gene groups. This is especially valuable in the study of condensates, where distinguishing different assemblies and RNA functions is challenging using traditional approaches. By integrating RNA-proximity ligation datasets into network-based analyses, we provide an approach to uncover new insights into the features organising specific RNA networks, aiding our understanding of higher-order transcriptome assemblage.

Overall, this study opens a new avenue for understanding the complex interplay between RNA identity and protein condensation potential in cellular organization and function, positioning smOOPs as potential players in the regulation of condensation processes.

### Limitations of the study

This study presents a new class of semi-extractable RNAs with high RBP occupancy determined by OOPS and global iCLIP. Both assays rely on UV crosslinking which has a strong bias towards uridines. Therefore, we cannot determine to what extent the observed higher RBP occupancy, especially in the A/U-rich cluster, reflects actual biological interactions. Although we performed orthogonal experiments to provide evidence that smOOPs are condensation-prone, they were not directly validated to undergo phase-separation processes. smOOPs RRI networks obtained by RIC-seq rely on pairwise interactions that show individual smOOPs can be near each other, though not necessarily all at the same time. Furthermore, it remains unclear whether their non-random association results from RNA condensation processes (such as co-assembly) or if it’s influenced by other factors. GO term analysis suggests different functions of smOOPs-encoded proteins from C- and A/U-rich clusters, however functional implications in the context of development and condensate formation could be further explored.

## Methods

### Cell culture

Low-passage, wildtype mouse pluripotent stem cells IDG3.2 PSCs (129S8/B6 background, ^48^ and TGFP/+;Foxa2tagRFP/+ (G9) ^49^ were cultured feeder-free, on cell culture dishes (TPP) coated with 0.1% Gelatin (ES-006-B, Millipore). Naïve PSCs (nPSCs) were maintained in N2B27 medium composed of 1:1 Neurobasal (21103049) and DMEM-F12 (11320074) medium containing N2 (17502001) and B27 (17504001) supplements, 1% Glutamax (35050061), 1% nonessential amino acids (11140050) and 0.1 mM 2-mercaptoethanol (31350-010) (all Thermo Fisher Scientific), 12 ng/mL LIF (104278, Qkine), with additional use of small molecule inhibitors: for a condition commonly named 2iLIF, 1 μM MEK inhibitor PD0325901 (1408, Axon Medchem) and 3 μM GSK3 inhibitor CHIR99021 (SML1046, Sigma). nPSCs were fed every day and split every 2-3 days using Accutase (A6964, Sigma). To transit from naïve to primed pluripotency state, the G9 PSCs were accutased and seeded onto gelatine and FBS-coated (EmbryoMax® ES Cell Qualified FBS, ES-009-B, Merck) plates in N2B27 medium supplemented with 1,000 U/ml LIF (ESGRO ESG1107, Merck), 12 ng/mL bFGF (100-18B, Peprotech), 20 ng/mL Activin A (338-AC-050, R&D Systems), and 2 μM IWP2 (3533, Tocris). Once the nPSCs reached primed pluripotency state - pPSCs - they can be maintained indefinitely using the abovementioned FAI medium. For further expansion, pPSCs G9 cells were maintained in FAI medium on cell culture dishes (TPP) coated with Geltrex (A1413302, Gibco), fed every day and split every 2-3 days using Accutase (A6964, Sigma).

To generate differentiated primitive streak progenitors (dPSCs) pPSCs were grown in Wnt3a-differentiation medium (N2B27 medium supplemented with 20 ng/mL ActA, 12 ng/mL bFGF and 250 ng/mL Wnt3a) for 24 hours prior collecting the cells.

### Semi-extractability and OOPS assays

pPSC G9 (p9) were prepared for experiment one day prior harvesting by plating them on 10 cm plates (TPP), and Wnt3a-differentiation medium was added to pPSCs for 24 hours to generate differentiated primitive streak progenitors (dPSCs). For this experiment only, 5% FBS (ES-009-B, Merck) was added to gelatin during coating for nPSCs (p20). Cells were prepared as follows: for semi-extractability and standard TRIzol RNA-seq controls one confluent 6-well was used per replicate. After washing cells 2x with PBS, 1 mL TRIzol-LS was added to the plate and scraped, then transferred to a 1.5 mL Eppendorf tube. Samples were snap-frozen and stored at -80°C. The semi-extractability assay was performed according to the protocol from ^20^: upon thawing and diluting TRIzol LS accordingly, the sample was heated for 10 minutes at 55°C and then sheared 40x through a 20G needle (the heating and shearing steps were omitted for control samples). Next, chloroform was added and mixed vigorously before the tube was centrifuged at 12,000x g for 15 minutes at 4°C. The aqueous phase was then mixed with 1.5x volume of 100% ethanol, and the isolation was continued with the RNeasy Plus Mini Kit (Qiagen) according to manufacturer’s instructions.

For OOPS, one confluent 10 cm plate was used per replicate, washed twice with ice-cold PBS before crosslinking them at 400 mJ/cm^2^ (Crosslinker CL-3000 at 254 nm (AnalytikJena)). After removing residual PBS, 1 mL TRIzol-LS was added to the plate before scraping off the cells content and transferring it to a 1.5 mL Eppendorf tube that was stored at -80°C. For RNA isolation, we followed the protocol provided by the Lilley Lab^50^. Briefly, the content of TRIzol LS tubes was first diluted with 1x volume nuclease-free water, then we added 1/5th volume of chloroform, mixed vigorously and centrifuged at 12,000x g for 15 minutes at 4°C. The aqueous and organic phase were carefully removed and the remaining interphase resuspended in fresh 1 mL TRIzol. Again, 200 µL chloroform was added before centrifugation and removal of aqueous and organic phases. Finally, the interphase was resuspended in 1 mL TRIzol once more before adding chloroform and separating the phases by centrifugation (12,000x g, 15 minutes, 4°C). This time, we removed as much of the aqueous and organic phases as possible and precipitated the remaining interphase (100 µL) using 100% Methanol (1 mL). After vortexing, the sample was centrifuged (14,000x g for 10 minutes at room temperature), supernatant was removed and the pellet digested using proteinase K (sro-3115828001, Roche) for two hours at 50°C. RNA was finally extracted by addition of 1 volume of phenol:chloroform (P3803, Sigma) and vortexing before centrifugation at 10,000x g for 10 minutes at room temperature. The aqueous phase was collected and RNA was precipitated for 20 minutes at room temperature with isopropanol. After centrifugation (12,000xg, 10 min, 4°C) the RNA pellet was washed twice using 75% ethanol and finally resuspended in nuclease-free water (AM9930, Thermo Fisher). All RNA samples were treated with Turbo DNase (AM2238, Thermo Fisher) for 30 minutes at 37°C to remove DNA contaminants and bead-purified using Agencourt AMPure XP (A63881, Beckman Coulter). Sequencing libraries were prepared using CORALL Total RNA-Seq V1 with RiboCop rRNA depletion (Lexogen) starting with 500 ng DNase-treated RNA as input. The resulting cDNA libraries were sequenced as single-end 100 bp reads on HiSeq at the Advanced Sequencing Facility at The Francis Crick Institute.

### Global iCLIP

iCLIP was performed according to the iiCLIP protocol^35^. First, 10 cm plates of nPSCs (G9, passage 17) and pPSCs (G9, passage 20) containing ∼1.5 mg of total protein mass per sample were lysed. Three replicates were prepared for each developmental stage. Briefly, cells were lysed upon UV-crosslinking (150 mJ/cm^2^ in Crosslinker CL-3000 at 254 nm (AnalytikJena)) and treated with RNase I before binding the RNA to an IR dye-labelled adaptor and separating the RNA-protein complexes on SDS-PAGE. After cutting the desired part of the membrane (from 40 kDa upwards), we digested the proteins and extracted the protein-bound RNA to prepare libraries for next-generation-sequencing (see Table S4 for adapter and primer sequences). Libraries were sequenced as paired-end 150 bp reads on NovaSeq at Clinical Institute of Special Laboratory Diagnostics, University Children’s Hospital at Ljubljana University Medical Centre.

### RIC-seq

Experiments were performed using nPSCs (G9, passage 20) and pPSCs (G9, passage 13)) in triplicates following the protocol by^51^) with some modifications. Two 10 cm confluent plates per sample were crosslinked using freshly prepared 0.5 mg/mL DSS (21655, Thermo Fisher Scientific) at room temperature for 30 minutes (rotating at 20 rpm). The reaction was quenched using 20 mM Tris (pH7.5) for 15 minutes at room temperature (rotating at 20 rpm). All centrifugation steps were carried out at 3500 rpm for 5 minutes at 4°C. After pelleting the cells, they were washed with PBS and then permeabilized (10 mM Tris-HCl (pH 7.5), 10 mM NaCl, 0.5 % Igepal, 0.3 % Triton X-100 and 0.1% Tween-20) for 15 minutes on ice. Cells were washed three times with 1X PNK buffer before fragmenting the RNA using 6U MNase (EN0181, ThermoFisher) at 37°C for 10 minutes. The following steps including FastAP treatment (EF0651, Thermo Fisher), pCp-biotin ligation (20160, Thermo Fisher), second FastAP and PNK treatment (M0201L, NEB) were performed as in the original protocol. Proximity ligation was performed using a different T4 RNA ligase (M0204L, NEB) and hence 1 mM ATP was added to the overnight reaction. Finally, cells were lysed using 200 μL proteinase K buffer (10 mM Tris-HCl pH 7.4, 100 mM NaCl, 1mM EDTA, 0,2% SDS) and 50 μL proteinase K (sro-3115828001, Roche) and an addition of 1.5 μLTurbo DNase (AM2238, Thermo Fisher), incubating at 37°C for 30 minutes, then 50°C for 60 minutes before adding Trizol-LS and snap freezing the samples. Once thawed and brought to room temperature, the samples were heated for 10 minutes at 55°C before adding the chloroform and precipitating RNA from the aqueous phase with isopropanol. The RNA was treated with Turbo DNAse once more and cleaned up using phenol:chloroform extraction as above. 21 μg of RNA was fragmented using 5x First Strand Synthesis Buffer (SuperScript IV, 18090050, Invitrogen) for 3.5 minutes at 94°C, immediately placed on ice and mixed with the MyOne Streptavidin C1 beads (65001, Invitrogen) to pull down biotinylated RNA (30 minutes at room temperature). Eluted RNA (10 μL) was extracted using phenol:chloroform method (see above). The biotinylation and pulldown efficiency were confirmed using dot blot assay before preparing sequencing libraries.

The biotin-enriched eluate was next subjected to 3’end dephosphorylation using PNK (M0201L, NEB) and FastAP (EF0654, Thermo Fisher) and purified using Agencourt AMPure XP beads (A63881, Beckman Coulter) before 3’end adapter ligation overnight at 20°C (Table S4). Once again, RNA cleanup was done with Agencourt AMPure XP beads before adapter removal (using Deadenylase (M0331, NEB) and RecJf exonuclease (M0264S, NEB)). Reverse transcription was performed according to the Superscript IV RT kit (18090050, Invitrogen) manual with the use of a custom RT primer (Table S4). Following cDNA cleanup (Agencourt AMPure XP beads) 5’ cDNA adapter was ligated using T4 DNA ligase (EL001, Thermo Fisher, without ATP) (Table S4). Samples were loaded onto 6% TBE-Urea gel (EC6865BOX, Thermo Fisher) and cDNAs exceeding 200 nt were excised from the gel and extracted using Crush-Soak Gel buffer (as per iiCLIP protocol^35^), followed by phenol:chloroform extraction. Precipitated cDNAs were stored at -20°C before performing PCR using P5/P7 standard Illumina primers and Phusion HF master mix (NE-M0531L, NEB). Ribosomal RNA contaminants were removed from the final library using Ribocutter which utilises Cas9-guided rRNA depletion^52^; for this, 275 gRNAs were designed against mature 5S, 18S and 28S rRNAs (obtained as 50 pmol oPool from IDT) (Table S4). Final libraries (∼10 nM) were treated with 4 μM sgRNAs for 30 minutes at 37°C and after beads purification, the libraries were reamplified with additional 6 cycles. They were sequenced as paired-end 150 bp reads on NovaSeq at Clinical Institute of Special Laboratory Diagnostics, University Children’s Hospital at Ljubljana University Medical Centre.

### HCR-FISH

Hybridization-chain reaction FISH was prepared by following the protocol from^36^ with slight modifications. The probes (8 pairs of probes per target, Table S4) were designed using https://github.com/rwnull/insitu_probe_generator^53^ against mature mRNAs, however not targeting exon-exon junctions (average expression for controls 23.6 +/- 16 TPM, for smOOPs 37.9 TPM +/- 31 TPM in semi-extractability total RNA-seq library). For imaging purposes WT IDG3.2 nPSCs were plated on Geltrex (A1413302, Gibco) coated 8-well glass-bottom ibidi plate (80827, ibidi) one day prior to fixation. After washing the cells with PBS, the cells were fixed by the fixation mixture (4% formaldehyde, 0.4% glyoxal, 0.1% methanol, 1x PBS). Amplification stage of a protocol to generate a tethered fluorescent amplification polymer lasted 10 hours. Cells were then washed as described in the protocol and finally mounted in 300 μl Fluoromount G (00-4958-02, Thermo Scientific). Images were acquired with a custom-built STED microscope (Abberior instruments) using a 1.2 NA 60× water immersion objective and lasers running at 80 MHz repetition rate. We excited fluorescently labelled mRNA by one of the three lasers at either 488, 561 or 640 nm, with 120 ps pulse length, with maximal power of 116 µW, 111 μW and 300 μW in the sample plane and DAPI stained nuclei with 405 nm laser with maximal power 810 mW in the sample plane. The laser powers used were 10% for 405 laser and 30% for the other three lasers. We acquired the fluorescence intensity using an avalanche photodiode with 500-550 nm, 580-625 nm or 650-720 nm filters (Semrock) in front. The combinations of lasers and detectors were as follows: 405 nm laser and 500-550 nm, 488 nm laser and 500-550 nm, 561 nm laser and 680-620 nm filter, and 640 nm laser and 650-720 nm filter. The dwell time in the pixel was 10 μs, the pixel size was set to 50 nm and the pinhole size was set to 1.07 AU to achieve a good confocal resolution.

### HCR-FISH image analysis

The image analysis was done in Fiji (Schindelin et al., 2012) using custom-made macros. Briefly, for each fluorophore background was subtracted (30.0 pixels rolling radius) and threshold was set accordingly to conform all different transcripts with the same fluorophore ((9,255) for channel for 561, (3,255) for the 488 channel and (33,255) for the 640 channel). Next, particle analysis was performed and all subsequent analysis was done using "Fiji’s particle analyser." Total intensity was calculated by multiplying mean intensity and area for each foci. The average of total intensity for the control transcripts in each fluorophore was used to normalise other values and obtain “Total intensity”.

### Reference annotation

For all analyses we used the GRCm39 build of the mouse genome with the <u>Gencode M27</u> annotation. We used a custom reference sequence built on this annotation for the alignment of hybrid reads, generated as previously described^54^. To unambiguously annotate the genes within hybrid reads, we used a flattened annotation produced by iCount-Mini (https://github.com/ulelab/icount-mini). Both are available for <u>download</u> with the Tosca pipeline^54^ (https://github.com/amchakra/tosca).

### RNA-seq data analysis

The sequencing reads were processed using nf-core/rnaseq version 3.4^55,56^ (https://nf-co.re/rnaseq/3.4). For differential expression analyses we used gene-summarised count tables generated by nf-core/rnaseq 3.4 (using Salmon) as input to DESeq2 version 1.44.0^57^. The design incorporated both stage (nPSCs, pPSCs, dPSCs) and assay (control, semi-extractability, OOPS) factors. Prior to running DESeq2, we pre-filtered the count matrix to retain only genes with at least 10 normalised counts in at least 6 samples. For each desired contrast, we extracted results using the Wald test and applied ashr Log2FoldChange (LFC) shrinkage^58^. To identify genes enriched in OOPS or semi-extractability compared to control, we selected genes with padj < 0.01 and LFC > 1 for each contrast (semi-extractability vs control or OOPS vs control at each stage). Genes passing these thresholds in both semi-extractability and OOPS were defined as smOOPs. For data visualisation across assays, we converted rlog-normalised count data into Z-scores for each stage and plotted it using the ComplexHeatmap package version 2.20.0.

### Global iCLIP data analysis

Sequencing reads were first demultiplexed using Ultraplex version 1.2.9 (https://github.com/ulelab/ultraplex) and then processed with the nf-core/clipseq version 1.0.0^59^ (https://nf-co.re/clipseq/1.0.0). We used BED files with crosslink positions and scores for all analyses. To normalise crosslinks by expression and correct for length for each transcript, we calculated crosslink density (CPM per kb) for exons and divided this by expression (semi-extractability TPM values obtained with Salmon).

### RIC-seq data processing

We trimmed sequencing adapters using Cutadapt^60^, then paired reads were merged with BBMerge^55^. The merged FASTQ files were used as input for Tosca v1.0.0^54^ to identify and analyse hybrid reads formed through RNA proximity ligation. To normalise RIC-seq gene-level counts to expression, we used TPM values calculated from RIC-seq nonhybrid reads summarised at the gene-level including all features (exons and introns) with featureCounts^61^.

### RIC-seq network inference and analysis

For each stage, we pooled the RIC-seq deduplicated hybrids files produced by Tosca, filtered out rRNA, tRNA and mitochondrial RNA containing hybrids, and retained the hybrid reads mapping to two different genes (representing inter-molecular interactions). We then summed the hybrid read counts corresponding to each gene pair to obtain gene-to-gene interaction frequencies. For each stage, we built a graph object, where genes were represented as nodes and edges were defined by the connections and their frequencies from the RIC-seq data, and analysed it using igraph^62,63^ version 2.0.3 in R 4.4.0. To assess the connectivity of the smOOPs subgraph, we compared the observed subgraph with control networks generated by degree-matched sub-sampling. For the degree-matched sampling method, we sub-sampled random subnetworks from the full RIC-seq network maintaining the same number of nodes as the number of smOOPs identified in the RIC-seq data at each stage and a similar degree distribution (allowing for +/-1 degree variation). We performed 10,000 iterations to build a null distribution for comparison. Connectivity metrics were calculated for each random subset and compared to those of the observed smOOPs subnetworks for nPSCs and pPSC separately. While we also considered degree-preserving randomization methods, they were deemed less suitable for this analysis due to the network topology and the high degrees of the smOOPs, which could bias the results by preserving inherent connectivity patterns, especially among high-degree nodes^64^. We also considered random sub-sampling, but found it overestimated smOOPs interconnectivity due to their high degree relative to most other nodes. In contrast, degree-matched sub-sampling, which directly compares smOOPs to randomly selected groups of nodes with similar degrees, provided a more appropriate baseline for comparison, and was used as control.

### PARIS data processing

We first trimmed sequencing adapters using Cutadapt and collapsed PCR duplicates with the readCollapse.pl script provided with the original publication^39^ (https://github.com/qczhang/icSHAPE). Subsequent processing was performed using Tosca v1.0.0^54^.

### Data preparation for deep learning

To define genes that do not exhibit the condensation prones features in all stages, we selected those with padj > 0.01 and |LFC| < 1.4 in each stage and then selected their intersection as the unified control set. To create a reference transcriptome for the smOOPs and control genes, we selected the most highly expressed transcripts, based on semi-extractability, for each of the three stages. For transcripts that varied across the stages, we selected the longest isoform among the most expressed transcripts at each stage.

To facilitate the training of the model on the selected datasets mapped across each transcript, we first encoded the data into a format suitable for deep learning. For both smOOPs and control transcripts, we extracted the genomic location of each exon from the reference genome. The RNA sequence for each exon was obtained based on their genomic coordinates using bedtools^65^, and served as the raw sequence input for the model.

Several nucleotide-resolution transcriptomic datasets were extracted and aligned to the exon genomic locations: global iCLIP crosslinks (this work); POSTAR3 database of CLIP binding peaks for 46 RBPs^41^ in mouse cell lines; Psoralen Analysis of RNA Interactions and Structures (PARIS)^39^; RNAfold^66^ predicting secondary RNA structures; and m^6^A methylation profiles^38^. To precisely assign scores to exon positions, we generated a list of values corresponding to the length of each exon, initialised to zero. The experimental data were mapped onto the exon’s genomic coordinates relative to its start position. Overlaps between experimental signals and exonic regions were located, and their scores were added to the list at the corresponding positions. The result was a list of experimental values corresponding to each position within the exon. For POSTAR3 peaks, each RBP peak was mapped in a binary manner, indicating only the presence or absence of a peak. After assigning scores, exons were concatenated back into transcripts, producing a list of feature scores spanning the entire transcript length for each feature. The global iCLIP, m^6^A modification sites and PARIS-Intra/Inter counts at each nucleotide were first normalised to transcript-level expression (TPM) estimated using the semi-extractability assay data, before encoding all feature layers of information over each smOOPs and control transcript.This approach preserves the positional importance and allows efficient feature extraction. Additionally, RNAfold was used to predict secondary RNA structures in dot-bracket notation, classifying each position as either single- or double-stranded., This classification was then encoded in binary form.

The datasets were randomly split into distinct training, validation, and testing subsets in a 70:15:15 ratio. Stratified sampling was employed to maintain consistent class proportions across subsets, and oversampling was applied to the minority group within each subset to balance class sizes. Each subset was stored independently and used directly for training, with dynamic data fetching and encoding implemented during the training process.

### Data encoding

During training, batches of data were procedurally encoded by iterating over the training dataset and processing the mapped information to a DL compatible array. For each batch, the transcript sequence information was first converted from its native nucleotide string format into a numerical representation. To ensure transparency and interpretability, we employed one-hot encoding for the sequence data, with each nucleotide represented as a binary vector: Adenine (A) as [1, 0, 0, 0], Cytosine (C) as [0, 1, 0, 0], Guanine (G) as [0, 0, 1, 0], and Thymine (T) as [0, 0, 0, 1]. Non-sequence features, such as global iCLIP crosslink scores, were encoded as one-dimensional arrays of floats. All encoded data layers, including the nucleotide sequences and other transcriptomic features, were stacked vertically for each transcript. To then standardise input lengths for neural network processing, sequences were padded with -1 to match the length of the globally longest transcript. The output class was similarly encoded using a one-hot encoding scheme, where class membership was represented by a binary vector. Such encoded transcripts were then stacked to a new dimension representing the batch and as such processed by the training function.

### Deep learning model architecture

The model architecture employs a previously optimised design based on^38^, combining multiple convolutional, recurrent, and fully connected layers (MLP) to effectively classify sequential data. This hybrid approach leverages the strengths of each layer type to capture diverse patterns in the input data, progressively transforming the sequences into representations suited for binary classification. Initially, we wanted to enable the architecture of the model to have the ability to achieve optimal classification performance, therefore a hyperparameter optimization process was conducted - defining the models architecture. Due to the complexity and size of the hyperparameter search space, a manual approach was impractical. Instead, a systematic search via Bayesian optimization, using the Optuna framework, was implemented, exploring batch sizes (8, 16, 32, 64), number of convolutional blocks (2 to 8), final number of CNN units (16 to 1024), CNN unit increase percentages (0.0 to 0.5), kernel sizes (3, 5, 7, 9), kernel size increases (1 to 4), dilation increases (1 to 4), dropout probabilities (0.0 to 0.4), L1-L2 regularisation values (0 to 0.01), max pooling size (2, 3, 4), GRU units (16 to 1024), Dense units (16 to 1024), learning rates (10^-5^ to 10^-3^), and normalisation methods (None, BatchNormalization, LayerNormalization). The optimisation was performed over 100 trials to identify the hyperparameter set that maximised AUROC on the validation set. The best-performing configuration was selected for final model training. The optimised model architecture starts with four convolutional blocks. Each block consists of a 1D convolutional layer with a progressively increasing number of filters of 108, 144, 192, and 256 respectively, and kernel size of 9. Dilation rates exponentially increase with each block, with the first block using a dilation of 1 and later blocks using 4, 16, and 64, respectively. Each convolutional layer is followed by layer normalisation, a ReLU activation function, and a Dropout layer with a rate of 0.34 to mitigate overfitting. Max-pooling (pool size of 4) is used after each convolutional block to reduce spatial dimensions and preserve key features. After the convolutional layers, a bidirectional GRU layer with 128 units per direction is employed to capture sequence dependencies, again followed by layer normalisation and dropout.

Following the GRU, two fully connected (Dense) layers are included, with 64 and 32 units, respectively, followed by layer normalisation and dropout. The final output layer consists of two units, with a softmax activation to predict binary classes. The Convolutional, GRU and Dense layers were configured with L1-L2 regularisation (λ = 1.2×10^-5^) to further mitigate overfitting in the learning process. The model contains between 1,149,986 and 1,203,446 trainable parameters, depending on the input shape. It was trained using the Adam optimizer, with a learning rate optimised to 10^-4^, and categorical cross-entropy as the loss function. The model was trained for unlimited epochs, employing early stopping with 20 epoch patience, based on validation AUROC, to stop training and return best weights when overfitting.

### The training of the powerset

To evaluate the predictive power of individual features and their combinations, we trained a separate model for each of the 127 subsets in the powerset of the 7 available datasets. For each subset, only the selected features were encoded as input to the model, ensuring that the model was trained specifically with that particular feature combination. Each model was trained in 8 replicates to account for variability and to ensure optimal performance for every subset. The trained models were saved after completion to enable further analysis of feature importance and interactions.

### Assessing model performance

The final performance of each of the 1016 trained models (127 subsets trained in 8 replicates) was established by predicting the combined validation and testing dataset, both of which were not previously seen by the model, ensuring robust validation and mitigating the optimisation overfitting. Of the 8 replicates trained on each subset, the best performing model, based on the AUROC, was selected, ensuring the robustness of the predictive power assessment. To identify the datasets that improved the predictive power of a model trained on any given subset of datasets, the difference in AUROC was calculated for all possible pairs of models where they were both trained on the same subset of datasets with the only difference being an additional inclusion of the unique dataset. This was further repeated for all possible pairs for each of the unique datasets and the differences were evaluated using a one-sample t-test to evaluate if the mean of differences is statistically different from zero, Using this technique, we could identify the datasets containing biological information that significantly improved the predictive capability of a model, irrespective of the number of other information layers included. To further assess the informational overlap between features, we conducted pairwise analyses of feature combinations. For each pair of features, we calculated AUROC scores for each subset of each item in their powerset, first by adding one feature from the pair, and then by adding both features. This approach allowed us to quantify the individual and combined contributions of each feature to model performance. To evaluate the extent of overlap, we calculated the difference between the maximum AUROC achieved when adding either feature individually and the AUROC obtained when both features were included. These differences were calculated across all subsets of the remaining five features, resulting in a comprehensive set of values for each feature pair. Averaging these differences provided insight into the distinct contributions of each feature combination with a one-sample t-test comparing the mean to zero.

### Explaining the trained weights

To evaluate the contribution of individual positions and feature components to the model’s predictions, we employed the integrated gradients (IG) method. The technique quantifies the importance of each nucleotide in the sequence by computing the gradient of the prediction output with respect to the input features, integrated along a path from a baseline of zeros to the actual input sequence. By summing these gradients across the entire path, IGs assign attribution/importance scores to each nucleotide, indicating its contribution to the prediction. We calculated the IGs for each of the models trained on individual datasets of sequence, global iCLIP, POSTAR3 peaks and PARIS-Intra. This approach enabled us to identify the regions within each transcript that had the greatest impact on the model’s classification decision, providing insights into which specific sequence elements were driving the condensation-prone smOOPs RNA predictions compared to controls.

### Gaining insight into the learned features

To gain insight into the global features the model has learned, not limited to a single example but integrated for the entire groups of transcripts, we firstly stacked the integrated gradients of each of the top 4 models to obtain the importance scores over all the predictive features for each transcript. These scores were grouped into 100 bins along the transcript length, and the average importance score within each bin was calculated to create a length-standardised distribution of importance across the sequence. We used Uniform Manifold Approximation and Projection (UMAP) to reduce the binned transcripts to 2D space, using correlation as the distance metric. Agglomerative clustering was then applied to group the transcripts into 2 clusters.

To elucidate the features defining each cluster we continued with the following analysis separately for each. We calculated the importance of nucleotide triplets across the transcripts. For each transcript, a sliding window approach was used to capture triplet sequences, and their integrated gradient scores were averaged for each triplet at every position. This reduced the transcript length by two. The triplet importance scores were then divided into 100 equal-length segments, and the scores were averaged across all transcripts. To further analyse the predictive importance of each POSTAR3 track, we calculated the average importance scores across all bins for each POSTAR3 RBP binding site in each transcript.

### Validation of the predictive importance by investigating original data

To assess nucleotide frequency differences between smOOPs and control transcripts in both clusters, we binned the sequences and averaged the presence of each nucleotide (A, C, G, T) across bins. For cluster 1, the sequence was separated into transcript regions, with the 5’UTR divided into 10 bins and the CDS and 3’UTR each divided into 50 bins. For cluster 2, the entire transcript was divided into 100 bins. The average nucleotide frequency within each bin was calculated separately for smOOPs and control transcripts. The differences in nucleotide frequency were determined by subtracting the control average from the smOOPs average for each nucleotide in each bin. To further investigate sequence composition, we calculated the frequency of each possible nucleotide triplet (3-mer) across the entire sequence for cluster 2 and within the transcript regions for cluster 1. The triplet occurrence frequency for control transcripts was subtracted from that of smOOPs transcripts for each cluster, yielding the difference in triplet usage. A similar approach was applied to the global iCLIP crosslinking signals. For cluster 1, the normalised signal was binned and averaged by transcript regions, with the 5’UTR divided into 10 bins and the CDS and 3’UTR into 50 bins each. For cluster 2, the signal was binned and averaged across the entire transcript into 100 bins. The median signal value for each bin was calculated for both smOOPs and control transcripts, and a 95% confidence interval for the median was estimated using bootstrapping with 1000 samples.

For cluster 1, we evaluated the impact of specific transcript regions on model predictions by masking either sequence, global iCLIP, POSTAR, and PARIS intramolecular interactions features. The entire region of the transcript was masked using zeros for 5’UTR, CDS, and 3’UTR individually. Models trained on each dataset were then used to predict outcomes based on these masked inputs. The AUROC curves were used to identify the region with the highest impact on the prediction when masked. For cluster 1 and 2, the number of PARIS intramolecular hybrids was calculated for each transcript part (5’UTR, CDS, 3’UTR) and full transcripts, respectively. These counts were normalised by transcript region length and expression levels.

### Investigation of the features of proteins encoded by smOOPs

To generalize the two-cluster annotation obtained in naïve PSCs (nPSCs) and apply it to all smOOPs transcripts, we trained an additional machine-learning (ML) model focused solely on RNA sequence features. As before, we used eight replicate training runs and retained the model achieving the highest area under the receiver operating characteristic curve (AUROC). We calculated the importance scores for each nucleotide position via integrated gradients, binned these scores, and clustered all smOOPs based on these aggregate importance profiles. We then evaluated how well the newly defined clusters (across all developmental stages) overlapped with the nPSC-specific clusters by measuring their percentage of intersection.

Next, to examine whether the two smOOPs clusters display distinct protein-level characteristics, we calculated amino acid frequencies for all proteins encoded by transcripts in each cluster and compared them against non-smOOPs. For each amino acid, we computed the log2 fold change in mean usage relative to non-smOOPs and determined statistical significance using Welch’s t-test.

We further assessed the extent of protein disorder in each cluster by extracting annotated intrinsically disordered regions (IDRs) from the UniProt Reviewed database^67^. For each protein, we recorded whether it contained any IDRs and calculated the average proportion of the protein length exhibiting disorder. Additionally, we used PICNIC^46^, a deep learning approach that leverages both sequence- and structure-derived information from AlphaFold2 models to evaluate how likely these proteins are to localize in biomolecular condensates.

Lastly, we performed gene ontology (GO) enrichment analysis to elucidate the functional distinctions of the two smOOPs clusters relative to the set of all expressed genes used as input to DESeq2. Using the STRING database^68^, we identified significantly enriched GO terms in each cluster across biological process, molecular function, and cellular component ontologies.

## Data and code availability

Newly produced and publicly available data were used for this work. Newly produced data was deposited on ArrayExpress under accession numbers E-MTAB-14762 (RNA-seq for semi-extractability and OOPS assays), E-MTAB-14763 (global iCLIP) and E-MTAB-14764 (RIC-seq). Raw PARIS data is available from GEO at GSE74353. Raw Ribo-seq data is available from GEO at GSE30839, with processed data (translation efficiency values) obtained from the supplementary files. SlamSeq metabolic RNAseq data was downloaded from GEO at GSE99978). Raw miCLIP (m^6^a) data is available at GEO at GSE169549. POSTAR3 CLIP datasets were obtained from the POSTAR3 platform (http://postar.ncrnalab.org/). The PICNIC scores for mouse proteome were obtained from the PICNIC platform (https://picnic.cd-code.org/). The code and notebooks to analyse the data and to produce the figures in this work are available at: https://github.com/ModicLab/smOOPs_project. The images and code used for HCR-FISH quantification and code are deposited at Zenodo (10.5281/zenodo.13860870).

## Supplemental information

Document S1. Figures S1–S5.

Table S1. Excel file containing identity of smOOPs and control transcripts, including cluster assignments related to Figures 1, 3, 4, and 5.

Table S2. Excel file containing the predictive performance of each trained model on validation dataset, related to Figures 3 and 5.

Table S3. Excel file containing the STRING GO enrichment analysis results, related to Figures 5.

Table S4. Excel file containing oligonucleotide sequences used in this study, related to Methods.

Data S1: Differential expression results in nPSCs between OOPS and control (standard TRIzol extraction).

Data S2: Differential expression results in nPSCs between semi-extractability and control (standard TRIzol extraction).

Data S3: Differential expression results in pPSCs between OOPS and control (normal TRIzol).

Data S4: Differential expression results in pPSCs between semi-extractability and control (standard TRIzol extraction).

Data S5: Differential expression results in dPSCs between OOPS and control (standard TRIzol extraction).

Data S6: Differential expression results in dPSCs between semi-extractability and control (standard TRIzol extraction).

## Acknowledgements

We would like to thank Flora Lee for establishing the RIC-seq library preparation method and Oscar Wilkins for designing guides for rRNA depletion using Ribocutter and helping with the Ultraplex demultiplexing tool. We are grateful to Charlotte Capitanchik for useful comments on the manuscript. We would like to thank Eneko Villanueva for providing us with a detailed protocol for OOPS, Yuanchao Xue for a detailed RIC-seq protocol and Joel Ryan for continuous advice on HCR-FISH imaging together with Petra Čotar and Luka Čehovin Zajc for automated image acquisition. We thank Thomas H Kapral and the other POSTAR3 authors for access to the full CLIPdb dataset. We also thank Tine Tesovnik and Jernej Kovač for their assistance and sequencing at the University Children’s Hospital at Ljubljana University Medical Centre. We also wish to acknowledge the Advanced Sequencing Facility and Nemo HPC at The Francis Crick Institute and HPC VEGA at the IZUM, the Institute of Information Science.

## Author contributions

T.K .: Conceptualization, Investigation, Project Administration, Formal Analysis, Writing - Original Draft Preparation, Writing - Review Editing; J.N.: Conceptualization, Methodology, Formal Analysis, Visualization, Data Curation, Writing - Original Draft Preparation, Writing - Review Editing; I.A.I.: Conceptualization, Methodology, Software, Formal Analysis, Visualization, Data Curation, Writing - Original Draft Preparation, Writing - Review Editing, Project Administration, Supervision; B.K.: Software, Formal Analysis; I.U.: Software; D.M.J.: Software, Formal Analysis; A.M.C.: Software, Writing - Review Editing, Supervision; N.M.L.: Resources, Funding Acquisition, Supervision; J.U.: Supervision, Resources, Funding Acquisition; M.M.: Conceptualization, Supervision, Project Administration, Resources, Funding Acquisition, Writing - Original Draft Preparation, Writing - Review Editing.

---

**Figure S1:**
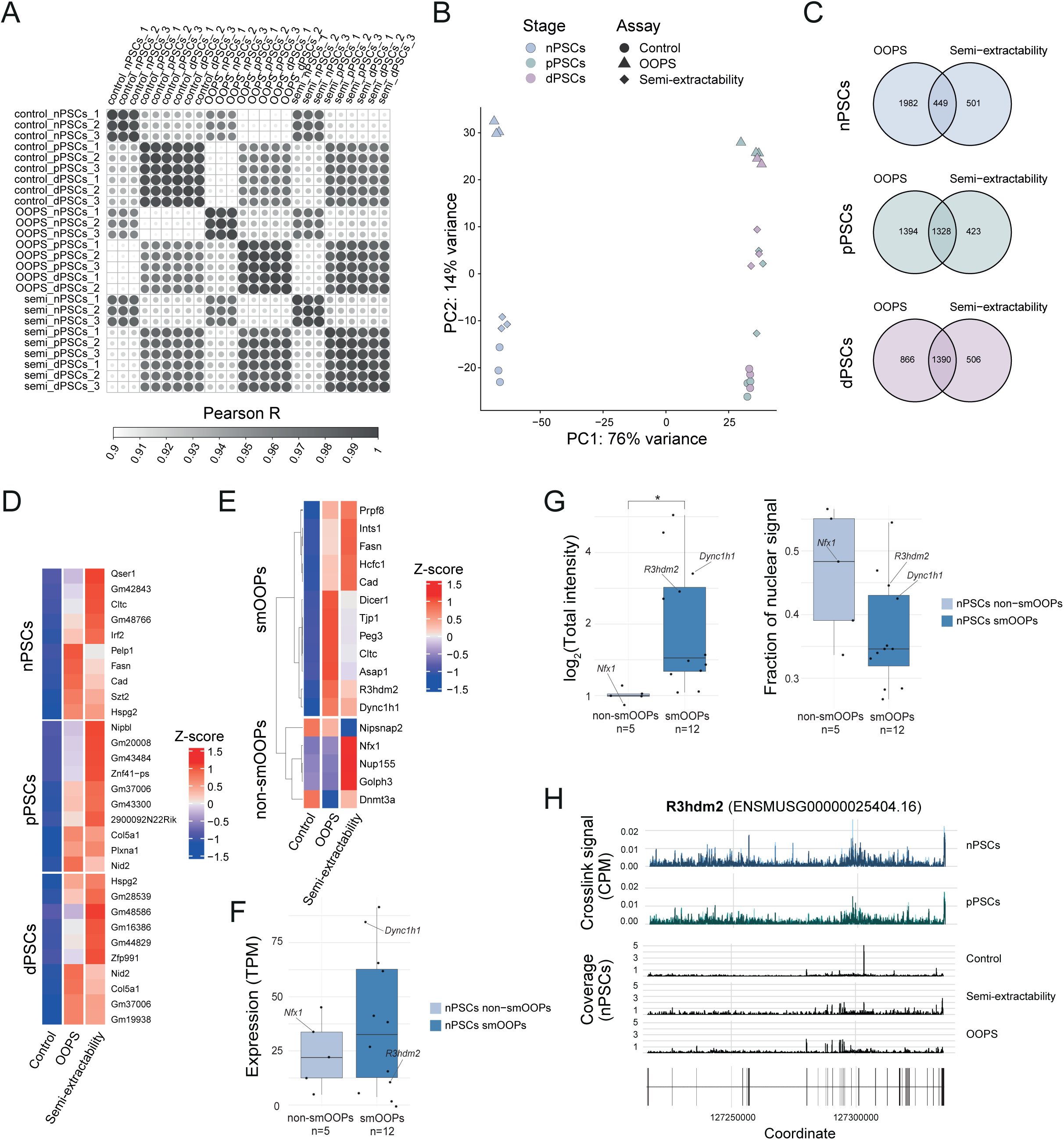
Identification of condensation-prone RNAs, related to Figure 1. A) Pairwise Pearson correlations between normalised gene-level counts for control, OOPS and semi-extractability assay samples in nPSCs, pPSCs and dPSCs shows high reproducibility among replicates. The dot size is proportional to the Pearson correlation coefficient. B) Principal component analysis using normalised gene-level counts using the top 2000 most variable genes separates samples on stage (PC1) and assay (PC2). C) smOOPs identification: Venn diagrams showing the overlap between semi-extractability assay and OOPS-enriched genes (differentially expressed vs control; LFC > 1, padj < 0.01) in nPSCs, pPSCs and dPSCs. D) Expression of example smOOPs for which fold changes in OOPS vs control are not equally high to the fold changes in semi-extractability assay vs control, and vice-versa. For each stage, the top five genes with highest fold changes in semi-extractability assay data and top five genes with highest fold changes in OOPS data are shown. E) Normalised expression profiles (Z-scores of rlog values) across assays in nPSCs for candidate mRNAs analysed by HCR-FISH. F) Distribution of expression levels in the semi-extractability assay data of transcripts tested using HCR-FISH F) Quantifications of the HCR-FISH data for smOOPs and non-smOOPs in nPSCs.. (Left) Boxplot showing total intensity (mean × area of each foci) normalised to the average intensity of control transcripts for each fluorophore used (Welch’s t-test, *p<0.05). (Right) Boxplot showing the fraction of foci for each transcript that are present in the nucleus. n indicates the number of different mRNAs against which the HCR-FISH probes were designed. G) Comparative visualisation of transcriptome data for an example smOOPs that was visualised by HCR-FISH, R3hdm2. Normalised crosslinking signal and coverage across assays was plotted using *clipplotr*^69^.

**Figure S2:**
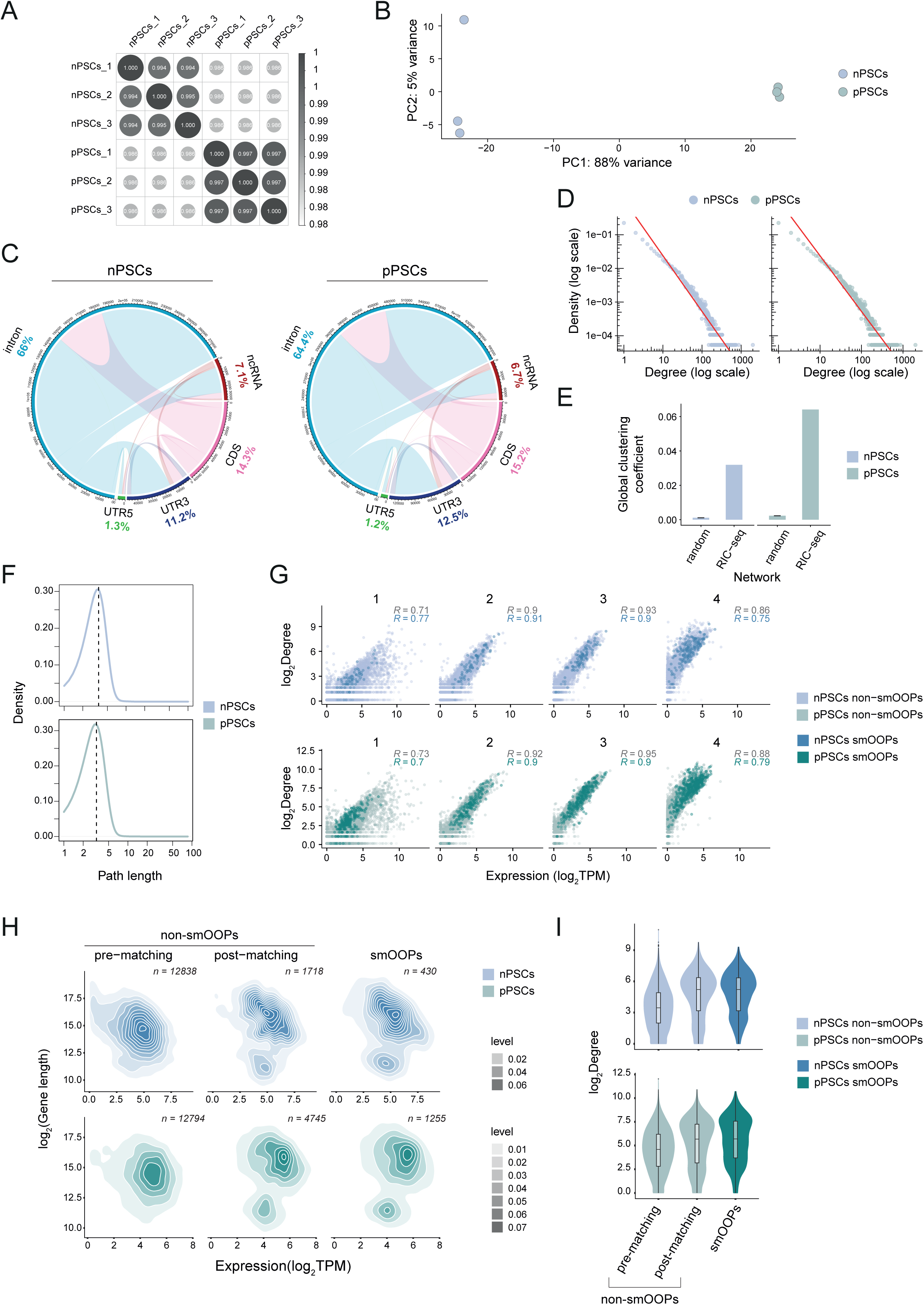
Characterisation of the RIC-seq data and networks in nPSCs and pPSCs, related to Figure 2. A) Pearson correlation between samples using normalised intermolecular gene-level hybrid read counts. B) PCA of the RIC-seq replicates in nPSCs and pPSCs based on the intermolecular hybrid reads using the top 2000 most variable genes. C) Circos plots showing the region links based on inter-molecular hybrid reads, with percentages indicating the proportion of interactions for each region type. D) Evaluation of the scale-free-like nature of the RIC-seq networks: degree distribution vs gene-counts on a log-log scale for the derived RRI networks. The red line indicates linear fit of the degree distribution on the log-log scale. E) Small world properties for the derived RRI networks: Mean global clustering coefficient compared to those of 100 random networks of the same size (equal number of nodes and edges). F) Small world properties for the derived RRI networks: Distribution of path lengths in the nPSCs and pPSCs RRI networks, with dashed vertical line indicating the average path length. G) Scatter plots showing the positive correlation between degree and expression across gene length quartiles (1-4). Blue and green Pearson coefficients (R) represent nPSCs and pPSCs smOOPs, respectively, while dark grey represents all genes. H) Quantile matching to select non-smOOPs with expression levels and gene lengths similar to smOOPs. The 2D density plots illustrate the distribution of all non-smOOPs, smOOPs, and expression- and length-matched non-smOOPs. I) Comparison of degree (number of distinct connections with other genes) distribution in the nPSCs and pPSCs RRI networks derived from RIC-seq between smOOPs, all non-smOOPs and the expression- and length-matched non-smOOPs, as identified in panel H.

**Figure S3:**
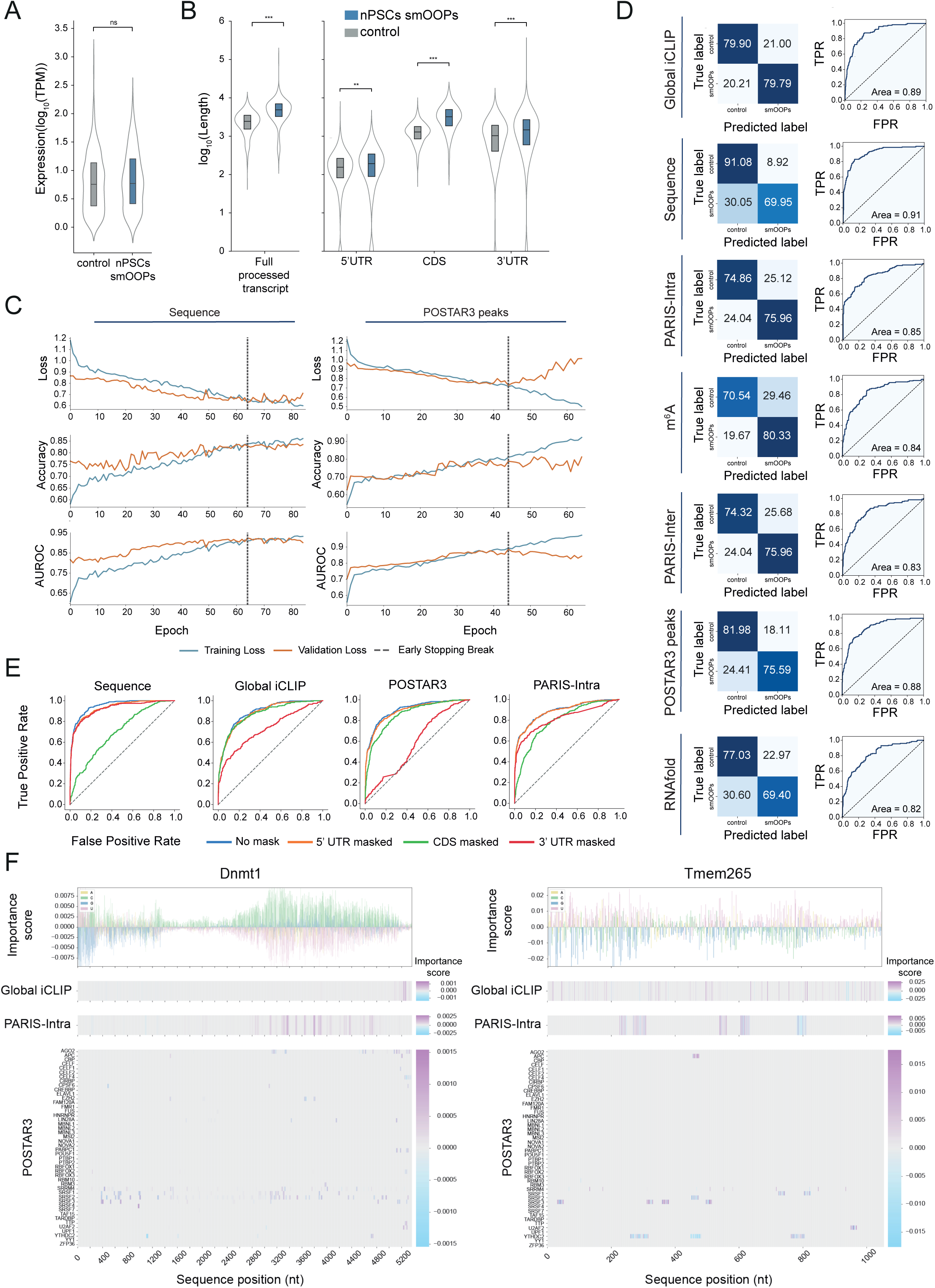
Training deep learning models to classify nPSCs smOOPs transcripts, related to Figure 3. A) Normalised expression levels for smOOPs and controls in the nPSCs semi-extractability assay data (Welch’s t-test, *p<0.05, **p<0.01, ***p<0.001). B) Distribution of full transcript lengths and transcript region-specific lengths for nPSCs smOOPs and controls (Welch’s t-test, *p<0.05, **p<0.01, ***p<0.001). . C) Training and validation loss, accuracy, and AUROC curves over training epochs for models trained on sequence (left) and POSTAR3 peaks (right) features. The dashed lines indicate the early stopping points based on validation AUROC performance. D) Classification performance of models trained on each individual feature set. (Left) Confusion matrices displaying the distribution of true and predicted labels for nPSCs smOOPs and control transcripts. (Right) AUROC curves that illustrate the true positive rate (TPR) versus the false positive rate (FPR), with the area under the curve (AUROC) indicated. E) AUROC curves showing the effect of masking specific transcript regions (5’UTR, CDS, 3’UTR) on the predictions of models trained solely on either sequence, global iCLIP, POSTAR, and PARIS-Intra features for cluster 1 smOOPs and control transcripts. F) Importance scores across transcript positions for *Dnmt1* (left) and *Tmem265* (right). Importance scores are derived from models trained on either global iCLIP, PARIS-Intra interactions, and the POSTAR3 peak dataset, shown in the respective tracks, with POSTAR3 RBP importance profiles shown for individual RBPs.

**Figure S4:**
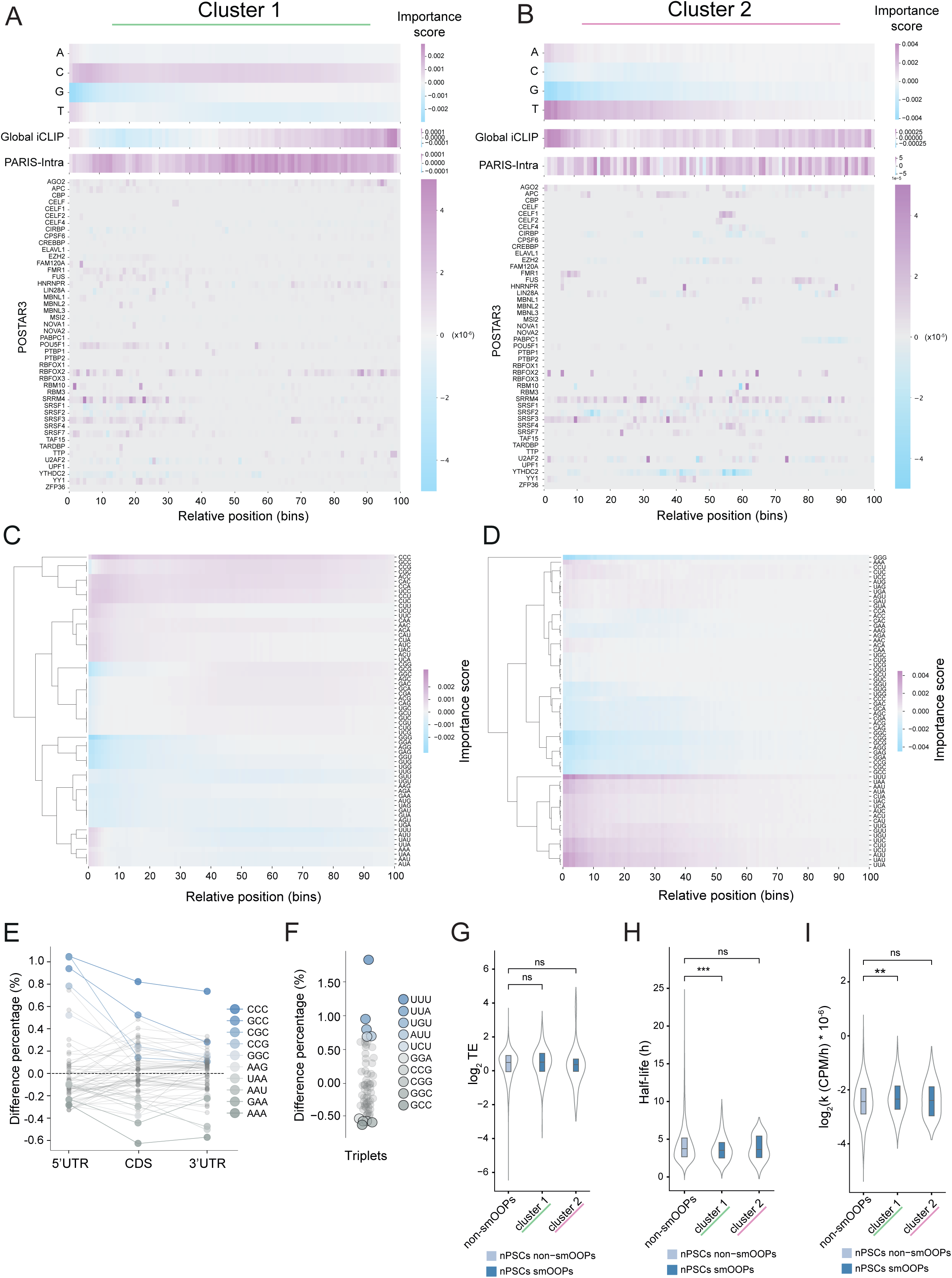
Identifying the positional importance of the most predictive features, related to Figure 4. A) Heatmap showing the average individual feature importance scores across all transcripts in cluster 1, divided into 100 bins over the entire transcript, with expanded individual POSTAR3 RBP importance. B) As in (A), but for cluster 2. C) Heatmap showing the importance scores of the sequence model averaged over a sliding window for individual triplets, binned into 100 bins over the entire transcript and averaged per bin for cluster 1. D) As in (C), but for cluster 2. E) The difference in average triplet content between cluster 1 nPSCs smOOPs and non-smOOPs for each transcript region. F) The difference in average triplet content between cluster 2 nPSCs smOOPs non-smOOPs. G) Translation efficiency in nPSCs for smOOPs and non-smOOPs mRNAs obtained by Ribo-seq^43^. H) Half-lives (h) of nPSCs smOOPs mRNAs compared to non-smOOPs, obtained by SLAM-seq^44^. I) Rate constants of decay (k in CPM/h) of nPSCs smOOPs mRNAs compared to non-smOOPs transcripts, obtained by SLAM-seq^44^.

**Figure S5:**
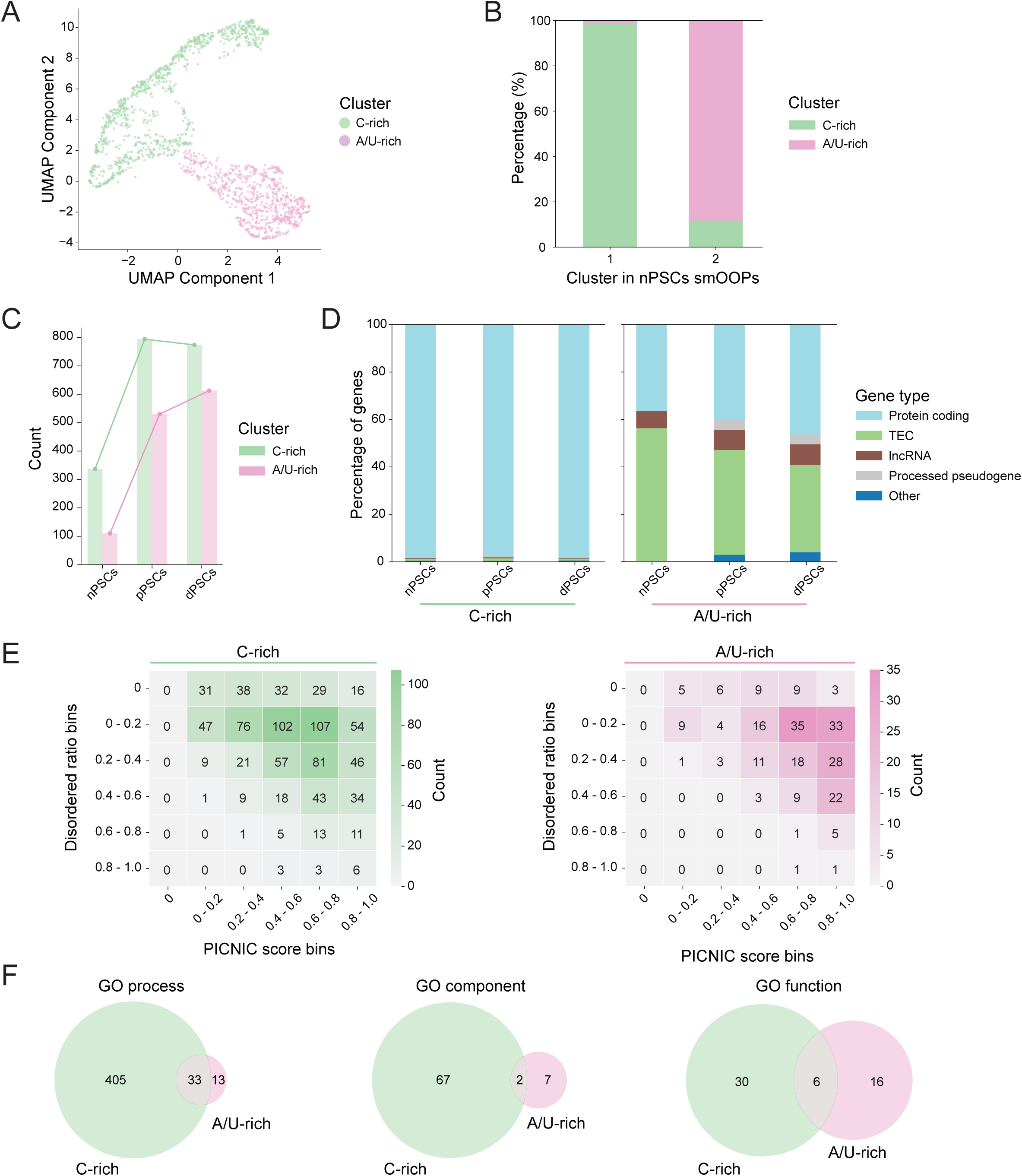
Functional and structural characteristics of smOOPs clusters, related to Figure 5. A) Scatter plot of UMAP components for all stages smOOPs transcripts based on binned importance scores for sequence only. Each dot represents a transcript, colour-coded by cluster. B) Barplot showing the overlap between clusters 1 and 2 of nPSCs smOOPs with the two clusters (C-rich, A/U-rich) obtained by clustering smOOPs from all developmental stages (nPSCs, pPSSc, dPSCs). C) Number of smOOPs transcripts in C-rich and A/U-rich clusters across developmental stages. D) Gene type composition of C-rich and A/U-rich smOOPs clusters across developmental stages. E) Relationship between protein disorder and PICNIC scores for proteins encoded by smOOPs in the C-rich and A/U-rich clusters. F) Distinct and overlapping GO term enrichments in C-rich and A/U-rich smOOPs clusters.

## References

1. Chen, X., and Mayr, C. (2022). A working model for condensate RNA-binding proteins as matchmakers for protein complex assembly. RNA 28, 76–87.

2. Forrest, K.M., and Gavis, E.R. (2003). Live imaging of endogenous RNA reveals a diffusion and entrapment mechanism for nanos mRNA localization in Drosophila. Curr. Biol. 13, 1159–1168.

3. Ramaswami, M., Taylor, J.P., and Parker, R. (2013). Altered ribostasis: RNA-protein granules in degenerative disorders. Cell 154, 727–736.

4. Boija, A., Klein, I.A., and Young, R.A. (2021). Biomolecular condensates and cancer. Cancer Cell 39, 174–192.

5. Modic, M., Grosch, M., Rot, G., Schirge, S., Lepko, T., Yamazaki, T., Lee, F.C.Y., Rusha, E., Shaposhnikov, D., Palo, M., et al. (2019). Cross-Regulation between TDP-43 and Paraspeckles Promotes Pluripotency-Differentiation Transition. Mol. Cell 74, 951–965.e13.

6. Alberti, S., and Hyman, A.A. (2021). Biomolecular condensates at the nexus of cellular stress, protein aggregation disease and ageing. Nat. Rev. Mol. Cell Biol. 22, 196–213.

7. Mittag, T., and Pappu, R.V. (2022). A conceptual framework for understanding phase separation and addressing open questions and challenges. Mol. Cell 82, 2201–2214.

8. Patel, A., Lee, H.O., Jawerth, L., Maharana, S., Jahnel, M., Hein, M.Y., Stoynov, S., Mahamid, J., Saha, S., Franzmann, T.M., et al. (2015). A Liquid-to-Solid Phase Transition of the ALS Protein FUS Accelerated by Disease Mutation. Cell 162, 1066–1077.

9. Wang, J., Choi, J.-M., Holehouse, A.S., Lee, H.O., Zhang, X., Jahnel, M., Maharana, S., Lemaitre, R., Pozniakovsky, A., Drechsel, D., et al. (2018). A Molecular Grammar Governing the Driving Forces for Phase Separation of Prion-like RNA Binding Proteins. Cell 174, 688–699.e16.

10. 10. Van Treeck, B., Protter, D.S.W., Matheny, T., Khong, A., Link, C.D., and Parker, R. (2018). RNA self-assembly contributes to stress granule formation and defining the stress granule transcriptome. Proc. Natl. Acad. Sci. U. S. A. 115, 2734–2739.

11. Cabral, S.E., Otis, J.P., and Mowry, K.L. (2022). Multivalent interactions with RNA drive recruitment and dynamics in biomolecular condensates in Xenopus oocytes. iScience 25, 104811.

12. Han, T.W., Portz, B., Young, R.A., Boija, A., and Klein, I.A. (2024). RNA and condensates: Disease implications and therapeutic opportunities. Cell Chem. Biol. 31, 1593–1609.

13. Jacq, A., Becquet, D., Guillen, S., Boyer, B., Bello-Goutierrez, M.-M., Franc, J.-L., and François-Bellan, A.-M. (2021). Direct RNA-RNA interaction between Neat1 and RNA targets, as a mechanism for RNAs paraspeckle retention. RNA Biol. 18, 2016–2027.

14. Trcek, T., Douglas, T.E., Grosch, M., Yin, Y., Eagle, W.V.I., Gavis, E.R., Shroff, H., Rothenberg, E., and Lehmann, R. (2020). Sequence-Independent Self-Assembly of Germ Granule mRNAs into Homotypic Clusters. Mol. Cell 78, 941–950.e12.

15. Maharana, S., Wang, J., Papadopoulos, D.K., Richter, D., Pozniakovsky, A., Poser, I., Bickle, M., Rizk, S., Guillén-Boixet, J., Franzmann, T.M., et al. (2018). RNA buffers the phase separation behavior of prion-like RNA binding proteins. Science 360, 918–921.

16. Parker, D.M., Tauber, D., and Parker, R. (2024). G3BP1 promotes intermolecular RNA-RNA interactions during RNA condensation. Mol. Cell. 10.1016/j.molcel.2024.11.012.

17. Trussina, I.R.E.A., Hartmann, A., Desroches Altamirano, C., Natarajan, J., Fischer, C.M., Aleksejczuk, M., Ausserwöger, H., Knowles, T.P.J., Schlierf, M., Franzmann, T.M., et al. (2024). G3BP-driven RNP granules promote inhibitory RNA-RNA interactions resolved by DDX3X to regulate mRNA translatability. Mol. Cell. 10.1016/j.molcel.2024.11.039.

18. Chen, X., Fansler, M.M., Janjoš, U., Ule, J., and Mayr, C. (2024). The FXR1 network acts as a signaling scaffold for actomyosin remodeling. Cell. 10.1016/j.cell.2024.07.015.

19. Wiedner, H.J., and Giudice, J. (2022). Author Correction: It’s not just a phase: function and characteristics of RNA-binding proteins in phase separation. Nat. Struct. Mol. Biol. 29, 1145.

20. Chujo, T., Yamazaki, T., Kawaguchi, T., Kurosaka, S., Takumi, T., Nakagawa, S., and Hirose, T. (2017). Unusual semi-extractability as a hallmark of nuclear body-associated architectural noncodingRNAs. EMBO J. 36, 1447–1462.

21. Zeng, C., Chujo, T., Hirose, T., and Hamada, M. (2023). Landscape of semi-extractable RNAs across five human cell lines. Nucleic Acids Res. 51, 7820–7831.

22. Queiroz, R.M.L., Smith, T., Villanueva, E., Marti-Solano, M., Monti, M., Pizzinga, M., Mirea, D.-M., Ramakrishna, M., Harvey, R.F., Dezi, V., et al. (2019). Comprehensive identification of RNA–protein interactions in any organism using orthogonal organic phase separation (OOPS). Nat. Biotechnol. 37, 169–178.

23. Srivastava, M., Srivastava, R., and Janga, S.C. (2021). Transcriptome-wide high-throughput mapping of protein–RNA occupancy profiles using POP-seq. Sci. Rep. 11, 1–15.

24. Ye, R., Zhao, H., Wang, X., and Xue, Y. (2024). Technological advancements in deciphering RNA-RNA interactions. Mol. Cell. 10.1016/j.molcel.2024.06.036.

25. Wu, T., Cheng, A.Y., Zhang, Y., Xu, J., Wu, J., Wen, L., Li, X., Liu, B., Dou, X., Wang, P., et al. (2024). KARR-seq reveals cellular higher-order RNA structures and RNA-RNA interactions. Nat. Biotechnol. 10.1038/s41587-023-02109-8.

26. Cai, Z., Cao, C., Ji, L., Ye, R., Wang, D., Xia, C., Wang, S., Du, Z., Hu, N., Yu, X., et al. (2020). RIC-seq for global in situ profiling of RNA–RNA spatial interactions. Nature 582, 432–437.

27. Zhao, H., Cai, Z., Rao, J., Wu, D., Ji, L., Ye, R., Wang, D., Chen, J., Cao, C., Hu, N., et al. (2024). SARS-CoV-2 RNA stabilizes host mRNAs to elicit immunopathogenesis. Mol. Cell 84, 490–505.e9.

28. Ying, Q.-L., Wray, J., Nichols, J., Batlle-Morera, L., Doble, B., Woodgett, J., Cohen, P., and Smith, A. (2008). The ground state of embryonic stem cell self-renewal. Nature 453, 519–523.

29. Kinoshita, M., Barber, M., Mansfield, W., Cui, Y., Spindlow, D., Stirparo, G.G., Dietmann, S., Nichols, J., and Smith, A. (2021). Capture of mouse and human stem cells with features of formative pluripotency. Cell Stem Cell 28, 453–471.e8.

30. 30. Kurek, D., Neagu, A., Tastemel, M., Tüysüz, N., Lehmann, J., van de Werken, H.J.G., Philipsen, S., van der Linden, R., Maas, A., van IJcken, W.F.J., et al. (2015). Endogenous WNT signals mediate BMP-induced and spontaneous differentiation of epiblast stem cells and human embryonic stem cells. Stem Cell Reports 4, 114–128.

31. Gouti, M., Tsakiridis, A., Wymeersch, F.J., Huang, Y., Kleinjung, J., Wilson, V., and Briscoe, J. (2014). In vitro generation of neuromesodermal progenitors reveals distinct roles for wnt signalling in the specification of spinal cord and paraxial mesoderm identity. PLoS Biol. 12, e1001937.

32. Clemson, C.M., Hutchinson, J.N., Sara, S.A., Ensminger, A.W., Fox, A.H., Chess, A., and Lawrence, J.B. (2009). An architectural role for a nuclear noncoding RNA: NEAT1 RNA is essential for the structure of paraspeckles. Mol. Cell 33, 717–726.

33. Pichon, X., Bastide, A., Safieddine, A., Chouaib, R., Samacoits, A., Basyuk, E., Peter, M., Mueller, F., and Bertrand, E. (2016). Visualization of single endogenous polysomes reveals the dynamics of translation in live human cells. J. Cell Biol. 214, 769–781.

34. Khong, A., and Parker, R. (2018). mRNP architecture in translating and stress conditions reveals an ordered pathway of mRNP compaction. J. Cell Biol. 217, 4124–4140.

35. Lee, F.C.Y., Chakrabarti, A.M., Hänel, H., Monzón-Casanova, E., Hallegger, M., Militti, C., Capraro, F., Sadée, C., Toolan-Kerr, P., Wilkins, O., et al. (2021). An improved iCLIP protocol. bioRxiv. 10.1101/2021.08.27.457890.

36. Choi, H.M.T., Schwarzkopf, M., Fornace, M.E., Acharya, A., Artavanis, G., Stegmaier, J., Cunha, A., and Pierce, N.A. (2018). Third-generation in situ hybridization chain reaction: multiplexed, quantitative, sensitive, versatile, robust. Development 145. 10.1242/dev.165753.

37. Ju, Y., Gong, J., Yang, Y.T., and Zhang, Q.C. (2018). Investigation of RNA-RNA interactions using the RISE database. Curr. Protoc. Bioinformatics 64, e58.

38. Modic, M., Kuret, K., Steinhauser, S., Faraway, R., van Genderen, E., Ruiz de Los Mozos, I., Novljan, J., Vičič, Ž., Lee, F.C.Y., ten Berge, D., et al. (2024). Poised PABP–RNA hubs implement signal-dependent mRNA decay in development. Nat. Struct. Mol. Biol., 1–9.

39. Lu, Z., Zhang, Q.C., Lee, B., Flynn, R.A., Smith, M.A., Robinson, J.T., Davidovich, C., Gooding, A.R., Goodrich, K.J., Mattick, J.S., et al. (2016). RNA Duplex Map in Living Cells Reveals Higher-Order Transcriptome Structure. Cell 165, 1267–1279.

40. Lorenz, R., Bernhart, S.H., Höner Zu Siederdissen, C., Tafer, H., Flamm, C., Stadler, P.F., and Hofacker, I.L. (2011). ViennaRNA Package 2.0. Algorithms Mol. Biol. 6, 26.

41. Zhao, W., Zhang, S., Zhu, Y., Xi, X., Bao, P., Ma, Z., Kapral, T.H., Chen, S., Zagrovic, B., Yang, Y.T., et al. (2022). POSTAR3: an updated platform for exploring post-transcriptional regulation coordinated by RNA-binding proteins. Nucleic Acids Res. 50, D287–D294.

42. Zhao, Y., Wang, J., Zeng, C., and Xiao, Y. (2018). Evaluation of RNA secondary structure prediction for both base-pairing and topology. Biophys. Rep. 4, 123–132.

43. Ingolia, N.T., Lareau, L.F., and Weissman, J.S. (2011). Ribosome profiling of mouse embryonic stem cells reveals the complexity and dynamics of mammalian proteomes. Cell 147, 789–802.

44. Herzog, V.A., Reichholf, B., Neumann, T., Rescheneder, P., Bhat, P., Burkard, T.R., Wlotzka, W., von Haeseler, A., Zuber, J., and Ameres, S.L. (2017). Thiol-linked alkylation of RNA to assess expression dynamics. Nat. Methods 14, 1198–1204.

45. Uversky, V.N. (2013). The alphabet of intrinsic disorder: II. Various roles of glutamic acid in ordered and intrinsically disordered proteins: II. Various roles of glutamic acid in ordered and intrinsically disordered proteins. Intrinsically Disord. Proteins 1, e24684.

46. Hadarovich, A., Singh, H.R., Ghosh, S., Scheremetjew, M., Rostam, N., Hyman, A.A., and Toth-Petroczy, A. (2024). PICNIC accurately predicts condensate-forming proteins regardless of their structural disorder across organisms. Nat. Commun. 15, 10668.

47. Faraway, R., Heaven, N.C., Digby, H., Wilkins, O.G., Chakrabarti, A.M., Iosub, I.A., Knez, L., Ameres, S.L., Plaschka, C., and Ule, J. (2023). Mutual homeostasis of charged proteins. bioRxiv. 10.1101/2023.08.21.554177.

48. Hitz, C., Wurst, W., and Kühn, R. (2007). Conditional brain-specific knockdown of MAPK using Cre/loxP regulated RNA interference. Nucleic Acids Res. 35, e90.

49. Scheibner, K., Schirge, S., Burtscher, I., Büttner, M., Sterr, M., Yang, D., Böttcher, A., Ansarullah, Irmler, M., Beckers, J., et al. (2021). Epithelial cell plasticity drives endoderm formation during gastrulation. Nat. Cell Biol. 23, 692–703.

50. Villanueva, E., Smith, T., Queiroz, R.M.L., Monti, M., Pizzinga, M., Elzek, M., Dezi, V., Harvey, R.F., Ramakrishna, M., Willis, A.E., et al. (2020). Efficient recovery of the RNA-bound proteome and protein-bound transcriptome using phase separation (OOPS). Nat. Protoc. 15, 2568–2588.

51. Cao, C., Cai, Z., Ye, R., Su, R., Hu, N., Zhao, H., and Xue, Y. (2021). Global in situ profiling of RNA-RNA spatial interactions with RIC-seq. Nat. Protoc. 16, 2916–2946.

52. Wilkins, O.G., and Ule, J. (2021). Ribocutter: Cas9-mediated rRNA depletion from multiplexed Ribo-seq libraries. 10.1101/2021.07.14.451473.

53. Kuehn, E., Clausen, D.S., Null, R.W., Metzger, B.M., Willis, A.D., and Özpolat, B.D. (2022). Segment number threshold determines juvenile onset of germline cluster expansion in Platynereis dumerilii. J. Exp. Zool. B Mol. Dev. Evol. 338, 225–240.

54. Chakrabarti, A.M., Iosub, I.A., Lee, F.C.Y., Ule, J., and Luscombe, N.M. (2023). A computationally-enhanced hiCLIP atlas reveals Staufen1-RNA binding features and links 3′ UTR structure to RNA metabolism. Nucleic Acids Res. 51, 3573–3589.

55. Bushnell, B., Rood, J., and Singer, E. (2017). BBMerge - Accurate paired shotgun read merging via overlap. PLoS One 12, e0185056.

56. Ewels, P.A., Peltzer, A., Fillinger, S., Patel, H., Alneberg, J., Wilm, A., Garcia, M.U., Di Tommaso, P., and Nahnsen, S. (2020). The nf-core framework for community-curated bioinformatics pipelines. Nat. Biotechnol. 38, 276–278.

57. Love, M.I., Huber, W., and Anders, S. (2014). Moderated estimation of fold change and dispersion for RNA-seq data with DESeq2. Genome Biol. 15, 550.

58. Stephens, M. (2017). False discovery rates: a new deal. Biostatistics 18, 275–294.

59. West, C., Capitanchik, C., Cheshire, C., Luscombe, N.M., Chakrabarti, A., and Ule, J. (2023). nf-core/clipseq - a robust Nextflow pipeline for comprehensive CLIP data analysis. Wellcome Open Res 8, 286.

60. Martin, M. (2011). Cutadapt removes adapter sequences from high-throughput sequencing reads. EMBnet.journal 17, 10–12.

61. Liao, Y., Smyth, G.K., and Shi, W. (2013). featureCounts: an efficient general purpose program for assigning sequence reads to genomic features. Bioinformatics 30, 923–930.

62. Csárdi, G., and Nepusz, T. (2006). The igraph software package for complex network research. In.

63. Csárdi, G., Nepusz, T., Müller, K., Horvát, S., Traag, V., Zanini, F., and Noom, D. (2024). igraph for R: R interface of the igraph library for graph theory and network analysis (Zenodo) 10.5281/ZENODO.7682609.

64. Horvát, S., and Modes, C.D. (2021). Connectedness matters: construction and exact random sampling of connected networks. J. Phys. Complex. 2, 015008.

65. Quinlan, A.R., and Hall, I.M. (2010). BEDTools: a flexible suite of utilities for comparing genomic features. Bioinformatics 26, 841–842.

66. Hofacker, I.L., Fontana, W., Stadler, P.F., Bonhoeffer, L.S., Tacker, M., and Schuster, P. (1994). Fast folding and comparison of RNA secondary structures. Monatsh. Chem. 125, 167–188.

67. UniProt Consortium (2025). UniProt: The universal protein knowledgebase in 2025. Nucleic Acids Res. 53, D609–D617.

68. Szklarczyk, D., Kirsch, R., Koutrouli, M., Nastou, K., Mehryary, F., Hachilif, R., Gable, A.L., Fang, T., Doncheva, N.T., Pyysalo, S., et al. (2023). The STRING database in 2023: protein-protein association networks and functional enrichment analyses for any sequenced genome of interest. Nucleic Acids Res. 51, D638–D646.

69. Chakrabarti, A.M., Capitanchik, C., Ule, J., and Luscombe, N.M. (2023). clipplotr-a comparative visualization and analysis tool for CLIP data. RNA 29, 715–723.

